# Mechanotransduction tuned by intermediate filaments promotes DNA repair in stiff environments

**DOI:** 10.64898/2026.10.30.763110

**Authors:** Cloé Dehours, Maëlle Locatelli, Cassandra Lepetit, Amanda Smelser, Hyunsu Lee, Julia Holmes, Qianqian Song, Mohamed Gaber, Sarvath Sanaullah, Maëlle Bertho, Sophie Lelièvre, Keith Bonin, Martin Guthold, Pierre-Alexandre Vidi

## Abstract

High mammographic density increases breast cancer risk. Dense tissues are rich in fibrillar collagen, forming a rigid extracellular matrix around the epithelium. Although cancer initiation and progression are closely tied to DNA damage and to the DNA damage response, if and how tissue stiffness affects genome maintenance is not fully elucidated. We found that repair factor recruitment is more efficient, and that DNA breaks are repaired faster by cells on rigid substrates. Mechanistically, hemidesmosomal integrins, intermediate filaments, the LINC complex, and YAP are key mediators of this mechanogenomic response. Rigid tissue contexts favor survival of cells with DNA damage, which may lead to the accumulation of somatic mutations, as identified here with RNAseq analyses of normal breast tissue. Our findings provide new insights into how tissue rigidity influences cancer risk.

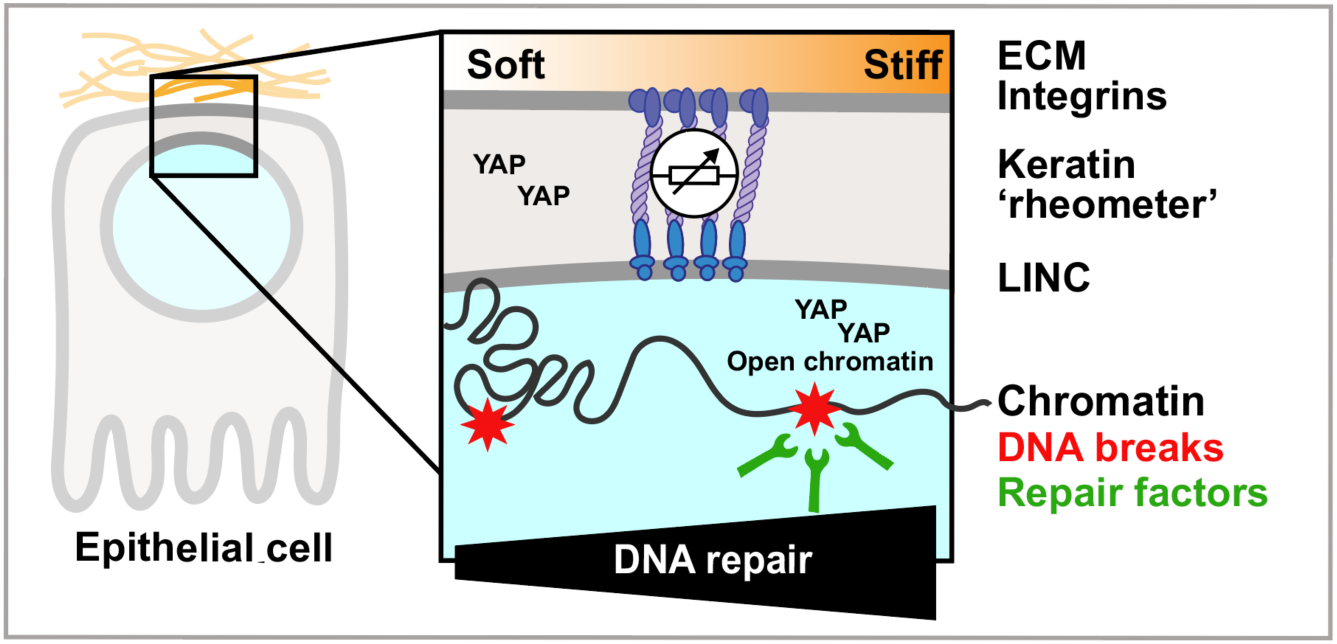

## Introduction

High mammographic density (MD) more than doubles the risk to develop breast cancer [1–4]. This association holds across ages [5] and breast cancer subtypes [6]. Radiodense breasts have a high proportion of extracellular matrix (ECM), in particular fibrillar collagen, and relatively less adipose tissue [7]. Breasts with high ECM content are stiffer than those with higher adiposity [8,9] and tissue stiffness *per se* is associated with breast cancer risk, even after adjusting for tissue density [10]. ECM stiffening is a hallmark of many solid tumors, including breast carcinoma. Whereas a stiff tumor microenvironment promotes cancer cell invasion and migration as a tumor progresses [11–16], the mechanisms linking tissue stiffness to cancer initiation are poorly understood.

Exposure to DNA damaging agents, as well as defects in the cellular responses to DNA damage are drivers of cancer risk in the breast and other organs [17]. Multiple breast cancer susceptibility genes are involved in DNA double-strand break (DSB) repair and reduced DNA repair capacity was measured in breast cancer patients compared to cancer-free controls [18,19]. Moreover, nonconservative, mutagenic, DNA repair pathways are more active in breast cancer patients and in women with familial breast cancer risk compared to the general population [20], indicating that repair accuracy is a molecular determinant of cancer risk. Evidence suggests a connection between breast density and genome integrity: increased histone H2AX phosphorylation (γH2AX), an epigenetic mark at DSBs, and shorter telomeres were measured in high vs. low MD breast tissues [21,22]. Yet, how stiff microenvironments within dense tissues may affect the DNA damage response (DDR) is not known.

Cells are influenced by the physical properties of their environment. Cell-ECM adhesion contacts are highly responsive to mechanical stimuli from the surrounding stroma [11,23,24] and a continuum of structural elements connects the ECM to the cell nucleus [25,26]. Cell membrane integrins, which bind ECM proteins and cytoskeletal elements, are mechanosensitive [27] and essential for differentiation, homeostasis, and cell survival [28–31]. Earlier work showed that alterations in cell-ECM contacts trigger genomic instability [32,33] and decrease DNA repair activity [34] in mammary epithelial cells. Intermediate filaments actively respond to mechanical cues in epithelial cells, contributing to cell stiffness adaptation [35,36] while transmitting forces to the nucleus and regulating nuclear mechanotransduction and deformation [36,37]. Although it is well established that physical cues from the cell microenvironment influence the chromatin state and gene expression programs [38–41], it is still unclear if and how these mechanical signals influence other genomic processes, in particular genome maintenance.

Here we show that the response to DNA damage is shaped by the stiffness of the cell microenvironment. In rigid contexts, the recruitment of DSB repair factors and the efficacy of DNA repair are enhanced, promoting cell survival after damage. We propose that in premalignant stiff environments, this elevated survival of damaged cells increases the mutational load, providing a mechanistic link between high MD and breast cancer risk.

## Materials and Methods

### Human tissue samples

Frozen needle biopsies of normal human breast tissues were obtained from the Susan G. Komen Tissue Bank (KTB; IU Simon Cancer Center, IN, USA). Collection of data and biospecimen by the KTB was approved by the IRB of Indiana University (Protocol #1011003097). Normal breast tissue from reduction mammoplasties was collected at the IU Health Arnett Hospital (Lafayette, IN). This tissue collection was approved by the IRB of Purdue University (protocol #1206012467). The fresh excision specimens were placed in RPMI, minced to 4-5 mm cubes, irradiated (3 Gy; Nordion Gammacell 220 irradiator), and left to recover for one or six hours in a cell culture incubator before freezing in optimal cutting temperature compound. Controls were mock-irradiated.

### Rheological characterization of human tissues

Atomic force microscopy (AFM) measurements were performed on frozen sections adhering to superfrost microscope slides. Tissues were thawed by immersing the slides 5 min in pre-warmed (30°C) PBS-PI [phosphate-buffered saline supplemented with 250 µM NaF, 10 µg/ml aprotinin (Gold Bio) and 1 mM phenylmethylsulfonyl fluoride], then stained 5 min with 4′,6-Diamidino-2-phenylindole dihydrochloride (DAPI; 0.5 µg/ml; Invitrogen) and washed with warm PBS-PI. Tissues were kept moist in this buffer during measurements. AFM measurements were taken with an MFP-3D-BIO system (Oxford Instruments) combined with an inverted microscope (IX73; Olympus). AFM microindentation and calculation of tissue elastic moduli was done as described [42]. We used a tipless, rectangular silicon nitride AFM cantilever (HYDRA2R-100NG-TL-10; nominal spring constant k = 0.011 N/m; AppNano) modified with a 5.3 µm diameter fluorescent melamine microsphere. The cantilever spring constant was determined using the ‘GetReal’ automated calibration function (thermal noise and Sader’s methods) of the AFM. The inverse optical lever sensitivity was determined directly by measuring the slope of the deflection voltage vs. z-distance curve on a hard glass surface. The stiffness (Young’s Modulus) of a selected sample region was determined using the built-in force map mode. In this mode, the AFM acquires 2D arrays of force distance curves, from which the stiffness of each pixel is calculated using the Hertz model of a hard sphere indenting a flat, elastic plane. The pixel size was 2 µm, and the area of the force map was between 60 µm x 60 µm and 80 µm x 80 µm. The indentation velocity of the force curves was between 2 and 5 μm/s. Regions of interest with breast epithelium were identified with phase contrast microscopy. A custom 3D-printed holder was used to match tissue slides to a gridded slide (Lovins Micro-Slide Field Finder, Electron Microscopy Science) to document the position of each structure (lobule/ducts) measured by AFM for subsequent confocal imaging (see below).

### Quantification of collagen alignment

Collagen alignment in human breast tissue was quantified based on collagen birefringence. Frozen breast tissue sections were thawed at room temperature and OCT (Optimal cutting temperature) compound was washed in distilled water. Tissues were fixed in formalin, rinsed in distilled water, and stained with picrosirius red F3BA (PSR) staining solution (Polysciences) for 90 minutes. Sections were briefly immersed in 0.1 N hydrochloric acid followed by 70% ethanol, then mounted with coverslips. Fibrillar collagen was imaged with polarized light microscopy at 20x magnification using an Olympus BX63 microscope. Collagen fiber organization was quantified using CT-FIRE and CurveAlign [43]. CT-FIRE was used to extract individual collagen fibers and CurveAlign to assess fiber alignment metrics at regions of interest (ROIs). For each acinus or duct, fiber alignment was measured in five ROIs (∼20 μm in diameter) located in the stroma directly adjacent to the structure to calculate average alignment scores.

### Cell culture and growth substrates

Non-immortalized post-stasis human mammary epithelial cells (HMECs; Lonza) were cultured in complete MammaryLife™ epithelial medium (Lifeline® Cell Technology) and passaged every 4-5 days up to passage 12. Immortalized breast epithelial cells from the HMT-3522 breast cancer progression series (S1, S2 and T4-2; a gift from Dr. M. Bissell) and MCF10A cells were propagated in H14 medium as described [44]. S1 cells expressing a dominant-negative β4-integrin truncated in its cytoplasmic tail [45] were obtained from Dr. Valerie Weaver. MCF10A cells expressing the C-terminal domain of 53BP1 fused to mCherry (mCh-53BP1_ct_) were generated by retroviral transduction of the mCherry-BP1-2 pLPC-Puro plasmid [46] (a gift from Dr. Titia de Lange; Addgene plasmid #19835). HepaRG liver epithelial cells were obtained from Biopredic Advancells and were cultured in William’s medium E supplemented with insulin (5 µg/ml), hydrocortisone hemisuccinate (5 µM), and 10% fetal bovine serum (FBS). U2OS cells stably expressing mCh-53BP1_ct_ [47] were cultured in DMEM/F12 supplemented with 10% FBS.

Hydrogels for 3D culture of S1 acini were prepared as described [48]. Briefly, collagen-I (4 mg/ml; Advanced Biomatrix, cat# 5201) was cross-linked with L-ribose (500 mM) for 10 days, then neutralized and mixed with reconstituted basement membrane (rBM; 20% final concentration; Culturex^TM^ Ultimatrix, Biotechne cat# BME001-05) and fibronectin (1 µg/ml final concentration; Sigma cat# F1141). A softer hydrogel was prepared in parallel with non-crosslinked collagen-I. Cells were mixed with the hydrogels (10^6^ cells/ml), seeded in chamber slides with a thin hydrogel coat, and cultured for 10 days.

Polyacrylamide (PA) gels (0.5 mm thick) were polymerized for one hour in silicone molds between coverslips. One coverslip was treated for gel adherence (1% 3-aminopropyltriethoxysilane followed by 0.5% glutaraldehyde) whereas the other coverslip was coated with Rain-X^®^ to facilitate removal after polymerization. The PA gels were made to target three distinct elastic moduli (200, 2000, and 12000 Pa) by adjusting the percentage of acrylamide (7.5, 5.5, and 3%) and of bis-acrylamide crosslinker (0.20, 0.15, and 0.08%), as described [49]. After rinsing three times with 200 mM HEPES, the top surface of the gels was activated by two applications of 1 mg/ml sulfo-SANPAH and 5 min exposures to UV (within 30 cm of a 30-watt germicidal fluorescent bulb). After three HEPES rinses, PA gels were coated with rBM (10% Matrigel®; Corning cat# 354234) for HMEC culture or with purified laminin (5 µg/ml; Gibco cat# 23017015) for S1 cell culture. For live-cell experiments, we used 35 mm dishes with thin (50 µm) PA gel islets with stiffnesses of 0.2, 2, and 12 kPa (Matrigen SoftIslet®), coated as described above. Silicone-coated 35 mm and 10 cm dishes (ExCellness) and 96-well imaging plates (Advanced Biomatrix) with defined stiffnesses were coated with laminin (20 µg/ml; Gibco) before cell seeding. These substrates were used when cells had to be retrieved for comet assay or western blot analyses, and for lentiviral transductions. HMEC cells were analyzed 12h after seeding on PA gels, whereas S1 and MCF10A cells were analyzed 10 days after seeding.

### Rheological characterization of growth substrates

Cell culture substrates were characterized by AFM and bead indentation measurements (Suppl. Fig. S14). PA gels were covered with culture medium and measured using an MFP-3D BIO AFM after one day of storage at 4°C. Tip-less HYDRA2R-100NG cantilevers were modified by gluing 5.3 µm diameter melamine formaldehyde fluorescent microspheres to the ends with Loctite Marine Epoxy, then coated in Rain-X® to reduce adhesion. The spring constant of the cantilever was determined using the Sader method in air, and the inverse optical lever sensitivity was determined using a deflection curve in culture medium [50]. The set point and trigger points were 0.5 V. Force curves were fit with the Hertz model, and the sample Poisson’s ratio was assumed to be 0.50 [51]. Each gel was measured at five different locations.

Elastic moduli were also determined at a larger scale by measuring the indentation depth due to gravity of stainless-steel beads (0.5 mm and 1 mm in diameter, DIT Holland B.V.). Indentation depth was imaged by embedding fluorescent beads (0.2 µm in diameter, Applied Microspheres) into the gel, then acquiring a z-stack underneath the steel bead using a LSM 880 confocal microscope. Depths were measured from maximal intensity projections in orthogonal views using ImageJ (https://imagej.nih.gov/ij/). Alternatively, substrates were coated with fluorescent beads and indentation depth was determined by reading focal positions on an Olympus IX83 microscope. Elastic moduli were calculated using the following formula:

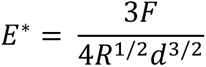

where *F* is gravitational force, *R* is the radius of the bead, and *d* is the indentation depth of the bead. Buoyancy was considered to be negligible because very little liquid was present on the surface of the gels.

### Gene silencing, recombinant protein expression, and chemical treatments

Transfection of S1 cells with nontargeting or plectin-targeting siRNAs (Dharmacon; OnTargetPlus), and with plasmids for expression of mCherry and mCherry-DN-KASH (Addgene plasmid #125553) was done with cell suspensions using Lipofectamine3000 (Invitrogen), according to the supplier’s recommendations. For depletion of keratins 8 and 18 in MCF10A, cultures in 96-well plates were transduced 48h after seeding with lentiviral particles encoding shRNA targeting *KRT8* (Dharmacon, #V3SH7590-227058229, sequence: ATGGACAACATGTTCGAGA) and *KRT18* (Dharmacon, #V3SH7590-226917880, sequence: AAGTAAAAGGCCTACAAGC) or a nontargeting control shRNA (Dharmacon, #VSC5985) at a concentration of 10^6^ TU/ml with polybrene (Sigma-Aldrich) at 10 µg/ml. Chemical treatments used in this study were colcemid, latrunculin A, okadaic acid (all from Cayman Chemical), paclitaxel (Accord), acrylamide (ThermoScientific), and verteporfin (Sigma). Concentrations and incubation times are indicated in figure legends.

### Induction of DNA damage to cells in culture

DNA damage, including DNA double-strand breaks, was induced with a ^137^Cs irradiator (JL Shepherd Mark I, Model 68A). For live cell imaging experiments, DNA breaks were induced using a miniature X-ray tube (Mini-X, Amptek) mounted on the microscope stage, at the maximal power setting (50 kV), and as described [52]. Dose rates were verified regularly using radiochromic films (Gafchromic^®^ EBT4, Ashland Inc). Unless indicated otherwise, a dose of 5 Gy was used, and cells were left to recover for 1h before analysis. Alternatively, DNA damage was chemically induced by treating cells with cisplatin (Accord) and bleomycin (TSZ Chemicals or Accord).

### Cell Survival Assays

For clonogenic assays, cells were cultured on PA gels, irradiated, and allowed to recover for 6 hours at 37°C before trypsinization and transfer to cluster plates (1,000 cells/well). After 5 days, cells were fixed (40% ethanol, 10% acetic acid) and stained with crystal violet. The plates were imaged using an Axio Zoom V16 microscope. Colonies, defined as groups of 5 cells or more, were assessed visually by an investigator blind to the treatment conditions. To assess viability by direct imaging of cells on PA gels, the LIVE/DEAD^®^ Viability/Cytotoxicity Kit (ThermoFischer) was used following the manufacturer’s protocol. Briefly, cells were stained with calcein-AM and ethidium homodimer-1 (30 min at 37°C), rinsed with phenol red-free culture medium, and imaged at 10x magnification with an IX83 Olympus microscope. Percentages of surviving cells were determined by quantifying the images with a custom ImageJ macro.

### Comet assay

Cells were placed on ice immediately after irradiation or after recovery, then trypsinized. The samples were resuspended in 0.5% low melting agarose and analyzed according to the manufacturer’s recommendations (Trevigen, neutral assay protocol). Comet images were taken with an IX83 Olympus microscope at 4x magnification and analyzed using OpenComet [53]. Median olive moments were calculated from 50-800 cells per condition and replicate.

### Immunostaining

Cells were fixed in 10% formalin (Merck cat# HT5011), permeabilized with TX-100, washed in PBS-glycine (50 mM), and incubated 2h in blocking buffer (10% goat serum in immunofluorescence buffer [IF; 130 mM NaCl, 13.2 mM Na_2_HPO_4_, 3.5 mM NaH_2_PO_4_, 0.1% bovine serum albumin, 0.05% NaN_3_, 0.2% Triton X-100, and 0.05% Tween 20]). Antibodies are listed in Table 1. They were diluted in blocking buffer and incubated on samples overnight at 4°C. Samples were washed three times with IF buffer, incubated with fluorescently labeled secondary antibodies (1h at room temperature), washed again with IF and stained with DAPI (0.5 µg/ml; 10 min). Samples were mounted with coverslips using ProLong Gold Antifade reagent (Invitrogen). Actin was detected with AF488-coupled phalloidin (Invitrogen; 1:400). Fluorescent signals were imaged using an Olympus IX83 epifluorescence microscope equipped with an ORCA-Quest qCMOS camera (Hamamatsu), using a 40x air objective (NA = 0.95). Excitation light was from an X-Cite TURBO LED illumination system (Excelitas). Emitted light was filtered using DAPI, GFP/FITC, mCherry/Cy3, and Cy5 filters (Chroma). For human tissues and 3D cell cultures, confocal images were taken with a Zeiss LSM880 confocal microscope using a 63x oil (NA = 1.4) objective.

**Table 1.** Antibodies.

| Antibodies | Source | Identifier | Concentration (application) |
|---|---|---|---|
| **53BP1** | Abcam | Ab36823 | 5 µg/ml (IS), 1 µg/ml (WB) |
| **α-tubulin (clone B-7)** | Santa Cruz | SC-5286 | 4 µg/ml (IS) |
| **α6-integrin (clone NKI-GoH3)** | Millipore | MAB1378 | 5 µg/mL (IS) |
| **β4-integrin (clone 3E1)** | Millipore | MAB1964-I | 1.7 µg/ml (IS) |
| **γH2AX (clone JBW301)** | Millipore | 05-636 | 2 µg/ml (IS) |
| **CK8 (clone EP1628Y)** | Abcam | Ab53280 | 0.07 µg/ml (IS) |
| **CK18 (clone DC10)** | Cell Signaling | 4548T | 0.8 µg/ml (IS) |
| **H3K9ac (clone Y28)** | Abcam | Ab12179 | 1 µg/ml (IS) |
| **H3K9me3 (clone D4W1U)** | Cell Signaling | 13969 | 0.2 µg/ml (IS) |
| **Ki67** | Thermo Fisher Scientific | PA5-19462 | 1 µg/ml (IS) |
| **Lamin A/C** | Santa Cruz | SC-7292 | 4 µg/ml (IS), 0.1 µg/ml (WB) |
| **Lamin B1** | Abcam | Ab16048 | 5 µg/ml (IS, WB) |
| **Lamin B2** | Abcam | Ab151735 | 2 µg/ml (IS) |
| **MDC1** | Abcam | Ab50003 | 2.3 µg/ml (IS) |
| **Par3** | Millipore | 07-330 | 13 µg/ml (IS) |
| **p-ATM (clone 10H11.E12)** | Santa Cruz | SC-47739 | 4 µg/ml (IS) |
| **Plectin (clone 10F6)** | Santa Cruz | SC-33639 | 4 µg/ml (IS) |
| **YAP (clone D8H1X)** | Cell Signaling | 14074 | 0.04 µg/ml (IS) |
| **ZO-1 (clone ZO1-1A12)** | Invitrogen | 33-9100 | 2.5 µg/ml (IS) |
| **ATM** | Abcam | Ab17995 | 1 µg/ml (WB) |
| **H2B** | Abcam | Ab52484 | 1.1 µg/ml (WB) |
| **Goat anti-Rabbit IgG, AFF488** | Thermo Fisher Scientific | A-11034 | 4 µg/mL (IS) |
| **Goat anti-Mouse IgG, AF488** | Thermo Fisher Scientific | A-11029 | 4 µg/mL (IS) |
| **Goat anti-Rabbit IgG, AF568** | Thermo Fisher Scientific | A-11011 | 4 µg/mL (IS) |
| **Goat anti-Mouse IgG, AF568** | Thermo Fisher Scientific | A-11031 | 4 µg/mL (IS) |
| **Goat anti-Rabbit IgG, AF647** | Thermo Fisher Scientific | A-21245 | 4 µg/mL (IS) |
| **Goat anti-Mouse IgG, AF647** | Thermo Fisher Scientific | A-21236 | 4 µg/mL (IS) |

### Live cell imaging

Time-lapse movies were captured at 40x magnification using the Olympus IX83 microscope. Cells were kept at 37°C and 5% CO_2_ in a stage-top incubator (Tokai Hit). Multiple regions of the dishes, corresponding to different substrate stiffnesses, were imaged sequentially over time using the motorized stage of the microscope and the multiple image acquisition function of the CellSens software (Olympus). Focus was maintained with a z-drift compensation system (TruFocus). The 475 nm LED was used for illumination and mCherry signals were collected with the Cy3 filter (600/37 nm). Low levels of excitation light (50 ms exposure) avoided photobleaching and phototoxicity.

### Image analyses

DSB foci numbers were evaluated visually in acini cultures (100-300 nuclei per condition and replicate) and in HMEC (25-150 cell for each replicate and condition). Scoring data were confirmed by evaluations from a second investigator who was blind to the treatment conditions. DSB foci in human tissue sections were also assessed visually (20-90 cells per patient and conditions) and the concordance between independent scorers was high (Suppl. Fig. S12J). In cell lines cultured as monolayers, repair foci were quantified using a custom macro in FIJI (https://fiji.sc). Briefly, nuclei were segmented using Cellpose [54], and foci were detected with a Laplacian of Gaussian (LoG) detector. The number of foci per nucleus was then quantified (100-1000 nuclei per condition/replicate). For live-imaging experiments, foci counts and clustering events were quantified visually. DSB foci analyses were done on maximal intensity projections from z-stacks to capture all foci in cell nuclei.

Cell cycle distribution was determined based on DAPI staining by imaging cytometry [55]. Illumination correction was done at the time of image acquisition in CellSens. Images were taken at 10x magnification, nuclei were segmented with StarDist [56] and the integrated density of DAPI signals was measured in FIJI. Histograms of the signal distributions, generated in GraphPad Prism, were based on a minimum of 3000 cells (typically 10000 - 30000) per condition/replicate.

Chromatin compaction was assessed with the coefficient of variation (CV) of DAPI signals, as described [57]. Cell nuclei were segmented from images taken with a 60x silicone oil immersion objective (NA = 1.30). The standard deviation of DAPI intensity values (σ) and mean DAPI intensity values (µ) were measured in FIJI to calculate CV = σ/µ. To quantify histone modifications, integrated densities from immunostaining images were calculated in cell nuclei ROIs obtained with Cellpose segmentation (N ≅ 500 cells per condition and replicate).

### Western Blot

Cells cultured on silicone-coated substrates were harvested by trypsinization. Proteins were denatured with 2% SDS and used for western blot analysis. Samples with equal protein amounts (30 µg), as determined using the DC kit from BioRad, were resolved on SDS-PAGE gradient gels (4-20%; BioRad), transferred to a nitrocellulose membrane, blocked with 5% nonfat milk in TBS-Tween (0.05%), and immunoblotted. Antibodies are listed in Table 1. The Clarity^TM^ Western ECL substrate kit (Bio-Rad) was used for detection. Signals were captured with a Fusion Solo S imager (Vilber) and quantified using ImageJ.

### Genomic analyses

RNA seq data from normal human breast tissue was used for low/high MD categorization based on signature and to call mutations. Data were obtained from the Genotype-Tissue Expression (GTEx) Project, version 8 [58]. Gene-level expression matrices were downloaded and preprocessed using standard normalization procedures. Samples were classified into low and high MD groups based on a previously published MD-associated 25 gene signature [59] based on mean expression per sample, which serves as a surrogate for tissue stiffness. Differential gene expression analysis was performed between high- and low-MD groups using edgeR package [60] within the R software, with multiple-testing correction applied to control the false discovery rate. Genes showing significant differential expression were subjected to Reactome pathway enrichment analysis using clusterProfiler [61] R package to identify biological processes associated with MD status. Mutation calls were retrieved from the GTEx portal. They correspond to breast tissue-specific SNVs inferred from RNA-seq, generated using the MuTect pipeline.

### Statistical analysis

Statistical analyses were done using GraphPad Prism 10. The Shapiro-Wilk test was used to test for normality before computing either t-test or one-way ANOVA and post-hoc tests. Nonparametric tests were used if the data did not pass the normality test (at alpha = 0.05). Statistical tests are indicated in the figure legends. For comet assays, significant outliers (Grubbs test, α = 0.05) were excluded. All statistical tests were two-sided. Unless specified otherwise, individual dots in the graphs represent summary values from independent biological replicates and bar graphs represent mean ± SEM.

## Results

### Stiff matrices enhance cell responses to DNA double-strand breaks

Our previous observation that the DNA damage response in breast acini is regulated by cell-ECM interactions via integrins [34] that are mechanosensors prompted us to compare two contrasting culture conditions: monolayers of breast epithelial cells (HMT-3522 S1; ’S1’) on stiff ECM-coated glass and S1 cells forming acini on a soft reconstituted basement membrane. In both settings, cells growth-arrest and differentiate [62] (Suppl. Fig. S1A). Yet, cells on glass accumulated significantly more DSB repair foci in response to ionizing radiation (IR) and to the radiomimetic agent bleomycin compared to acinar cells (Suppl. Fig. S1B-C). Next, we compared S1 acini in soft ECM to acini grown in ECM cross-linked with ribose. The latter had significantly more IR-induced γH2AX and 53BP1 DSB foci (Fig. 1A). Rheology measurements showed a 60% increase in the elastic modulus of cross-linked gels (Fig. 1B). Hence, the mechanical properties of the cell microenvironment affect the response to DSBs.

**Figure 1.**
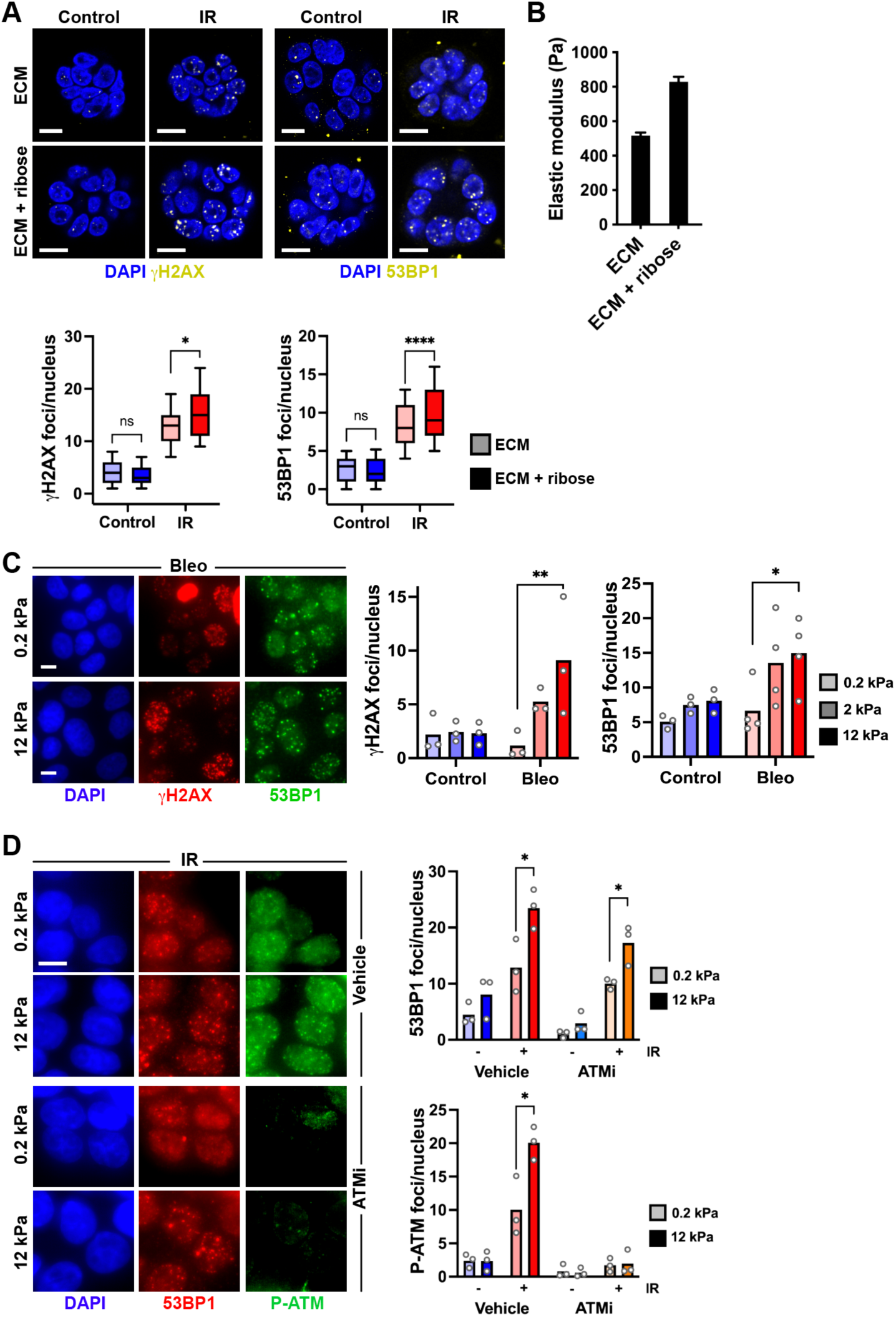
Extracellular matrix stiffness influences DSB repair foci accumulation in breast epithelial cells. (**A**) Formation of γH2AX and 53BP1 repair foci in S1 acini in soft ECM and in ECM stiffened with ribose, in response to ionizing radiation (IR). *, *P* < 0.05 and ****, *P* < 0.0001 (Kruskal-Wallis and Dunn’s multiple comparison test; N ≥ 200 nuclei from two independent experiments). (**B**) Elastic moduli of ECM used in (A), determined by bead indentation (N = 3). (**C**) γH2AX and 53BP1 repair foci in S1 cells on polyacrylamide gels with different elastic moduli. Cells were either untreated (control) or treated with bleomycin (Bleo; 20 mU/ml, 1h). Scale bars, 10 µm. *, *P* < 0.05 and **, *P* < 0.01 (ANOVA and Bonferroni’s multiple comparison test). (**D**) 53BP1 and P-ATM foci in S1 cells irradiated on substrates with different stiffnesses. Cells were treated with ATMi (Ku55933; 10 µM, 2h) as indicated. *, *P* < 0.05; ns, not significant (unpaired t-test).

To isolate effects driven by mechanical properties of the growth substrate from acinar morphogenesis, we used polyacrylamide gels with different elastic moduli. The stiffness of these gels can be tuned reproducibly [63] while keeping the concentration of ECM ligands constant. Gel stiffness ranged from 0.2 to 12 kPa, spanning values measured in the mammary gland [8,9,11,64]. The radiomimetic drug bleomycin induced higher levels of γH2AX and 53BP1 DSB repair foci on stiff vs soft gels (Fig. 1C). Similarly, S1 cells irradiated on a stiff substrate accumulated more 53BP1 foci than cells cultured in soft conditions (Fig. 1D). ATM autophosphorylation at serine 1981 (P-ATM) is an early DDR response stabilizing the active kinase at DSBs [65]. On stiff ECM, the number of IR-induced P-ATM foci was significantly higher than on soft ECM. Inhibiting ATM activation with ATMi blunted P-ATM signals and modestly reduced IR-induced 53BP1 foci but did not block the effect of substrate stiffness on the DDR (Fig. 1D). The DDR is influenced by cell cycle status. Yet, S1 cell cycle distribution (based on DNA content) and proliferation (based on Ki67 positivity) were similar across the different stiffnesses (Suppl. Fig. S2A-B). We also found that 53BP1 expression—which may affect our ability to identify repair foci—was not altered by the stiffness of the substrate (Suppl. Fig. S2C). Importantly, as shown and discussed below, the amount of DSBs generated by IR (measured with comet assay) was not influenced by the stiffness of the growth substrate. We obtained similar results with non-immortalized human mammary epithelial cells (HMEC). IR-induced γH2AX and 53BP1 foci counts were ∼50% higher in HMEC on stiff gels than on soft gels, and cells on matrices of intermediate stiffness had intermediate levels of IR-induced foci (Suppl. Fig. S3A). Substrate stiffness did not affect HMEC proliferation, nor the level of IR-induced DSBs (Suppl. Fig. S3B-D). The results indicate that, *in vitro*, the mechanical properties of the microenvironment affect the cellular response to DNA damage rather than DNA damage itself.

To determine if the influence of ECM stiffness on the DNA damage response is retained after malignant transformation, we analyzed invasive ductal carcinoma cells (HMT-3522 T4-2; ’T4-2’). Induction of DSB foci by IR on the soft (0.2 kPa) substrate was less pronounced than on stiffer (2 and 12 kPa) substrates (Suppl. Fig. S4A). Again, cell cycle distribution was not altered by substrate stiffness (Suppl. Fig. S4B) and levels of IR-induced DSBs measured by comet assay were similar across stiffness conditions (Suppl. Fig. S4C).

To assess generalizability to other organs, we analyzed the radiation response of liver epithelial cells (HepaRG). Since the liver is substantially stiffer than the normal breast epithelium, we extended the range of stiffnesses and included cells growing on glass. Higher accumulation of radiation-induced foci was observed on stiff substrates (12 kPa and glass) whereas radiation responses were similar for cells growing in softer (0.2 and 2 kPa) conditions (Suppl. Fig. S5A). Again, cell cycle distribution and IR-induced DSB levels were not different across stiffness conditions (Suppl. Fig. S6B-C). By contrast, U2OS osteosarcoma cells—derived from the inherently stiff bone tumor environment and known to be poorly mechanoresponsive—showed constant IR-induced 53BP1 foci numbers across stiffnesses (Suppl. Fig. S6). In summary, mammalian cells respond differently to DNA damage according to the stiffness of their growth substrate. This effect varies for cells of different origins and may reflect mechanogenomic responses tuned to physical contexts.

### ECM stiffness increases breast epithelial cell survival to DNA damage

These differences in DSB factor recruitment prompted us to assess cell survival after DNA damage as a function of ECM compliance. In clonogenic assays, HMEC irradiated on stiff (12 kPa) substrates displayed significantly higher survival than cells on softer (2 kPa) substrates (Fig. 2A). Cells grown on yet softer (0.2 kPa) substrates failed to form colonies after transfer to assay plates and could not be assessed. Using viability staining to avoid the confounding effect of transferring cells on plastic, we measured higher viability for S1 cells on stiff compared to soft substrates following exposure to genotoxic chemotherapy and paclitaxel (Fig. 2B-C), indicating pro-survival effects of stiff ECM not restricted to the DNA damage response.

**Figure 2.**
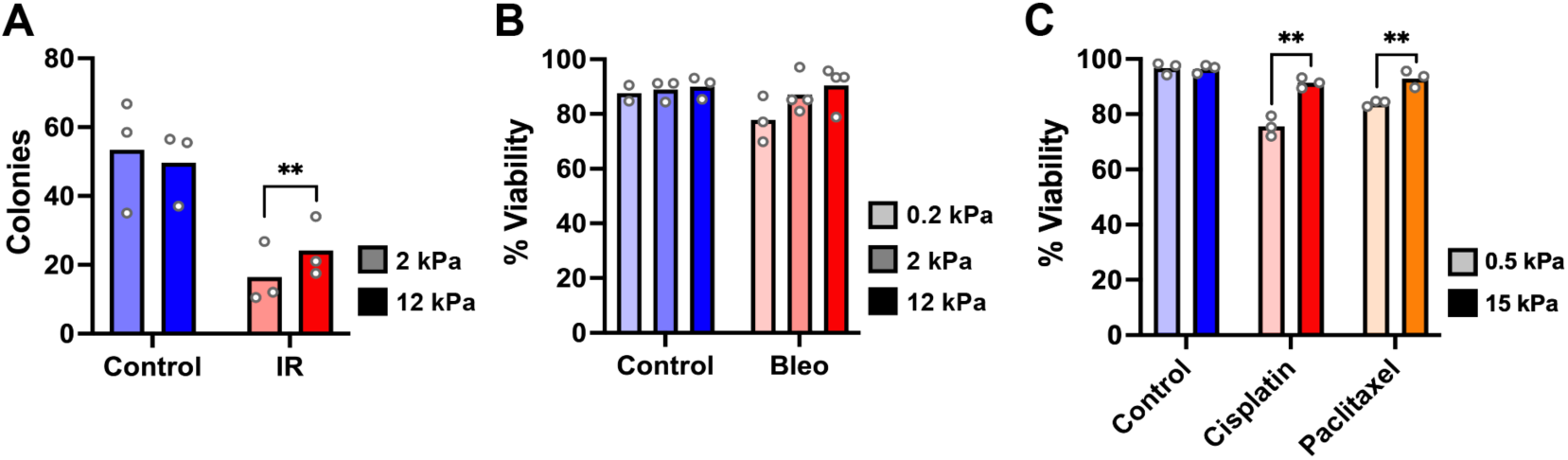
Stiff ECM increases cell survival to DNA damage. (**A**) Clonogenic assay with HMEC irradiated (IR) on substrates with different elastic moduli. Controls were mock-irradiated. (**B-C**) Viability of S1 cells on substrates with different stiffnesses, after treatments with bleomycin (20 mU/ml; 24h) (B) or cisplatin (100 µM; 48h) and paclitaxel (1 µM; 48h) (C). **, *P* < 0.01 (t-test).

### Cells on stiff matrices have faster DSB repair kinetics

We compared DSB repair by S1 cells on soft and stiff ECM with time course of comet assays. As mentioned above, IR-induced DNA damage levels were similar across stiffnesses, but DSBs persisted longer in the soft growth condition (Fig. 3A), suggesting faster repair on rigid ECM.

**Figure 3.**
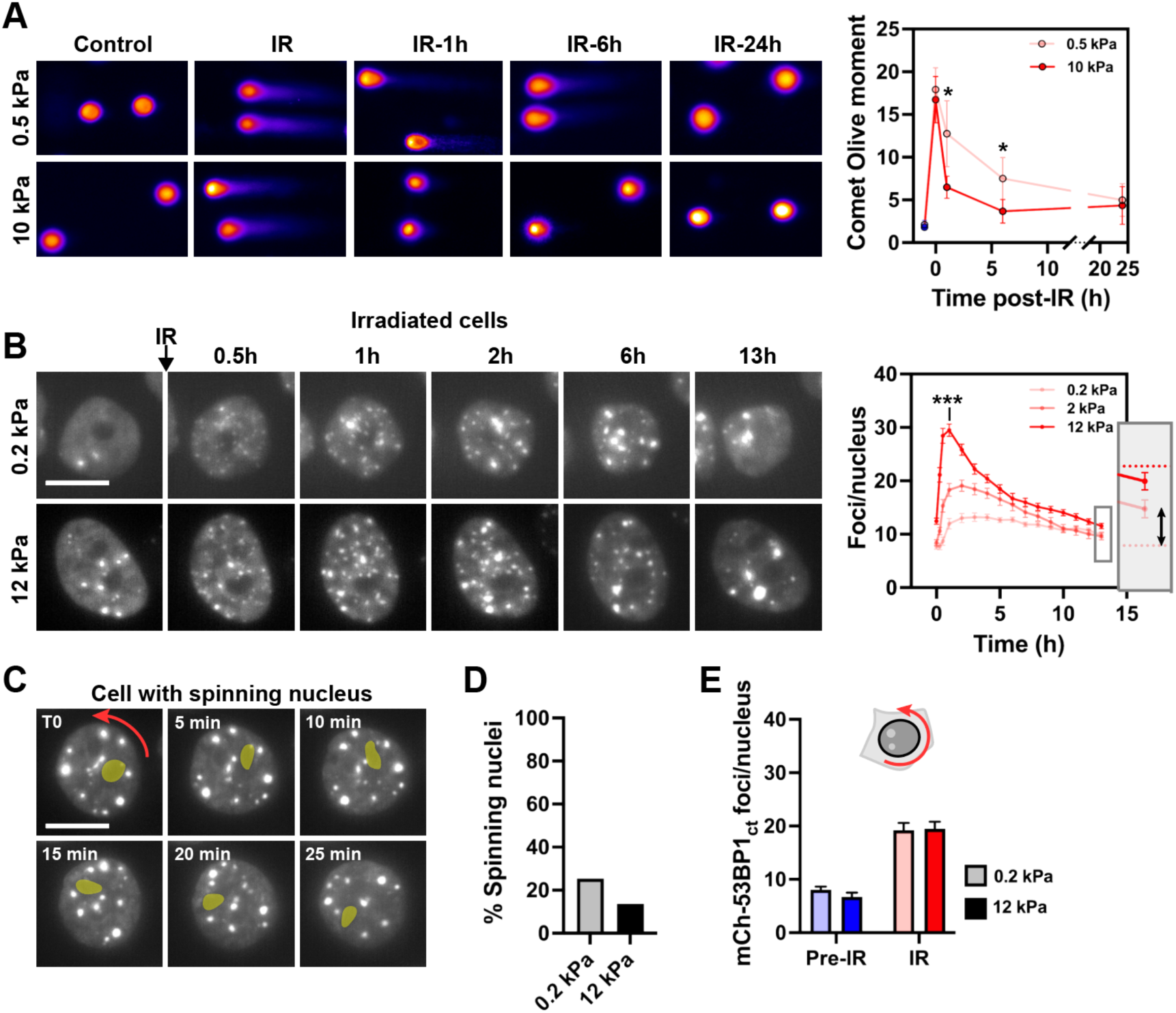
DSB repair kinetics depends on the mechanical properties of the ECM. (**A**) Neutral comet assay with S1 cells on soft or stiff substrates, performed at different time points after irradiation (IR). *, *P* ≤ 0.05 (N = 6; ratio paired t-test). (**B**) Representative mCherry-53BP1_ct_ signals and foci counts from cells on soft or stiff substrates following IR. ***, *P* = 0.0003 (ANOVA and Bonferroni’s multiple comparison test; 1h time point; N = 25-30 cells per condition). Dotted lines in the graph inset show average foci counts pre-irradiation (0.2 and 12 kPa conditions). (**C**) Illustration of a spinning cell nucleus (defined as ≥5 x 360° rotations over a 12h period). A nucleolus is highlighted (yellow) as reference. (**D**) Proportion of cells with spinning nuclei in soft and stiff contexts (N > 200 nuclei). (**E**) mCherry-53BP1_ct_ foci counts for cells with spinning nuclei, before and 2h after IR (N = 20-30 cells). Scale bars, 10 µm.

To visualize repair dynamics in real time, we combined live cell imaging with a miniature X-ray source [52] and used MCF10A breast epithelial cells expressing a DSB sensor. The sensor consists of the minimal foci forming region of 53BP1 tagged with mCherry (mCh-53BP1_ct_) [66,67]. As expected, mCherry-labeled foci overlapped with endogenous 53BP1 and γH2AX foci detected by immunostaining (Suppl. Fig. S7A). Cells were cultured in dishes with thin polyacrylamide disks with different elastic moduli, enabling simultaneous time-lapse imaging on different stiffnesses, before and after irradiation (Suppl. Fig. S7B). Again, MCF10A cell cycle distribution did not vary on the different stiffnesses (Suppl Fig. S7C). We also verified that live imaging was not inducing DNA breaks (Suppl. Fig. S7D, Suppl. Movie S1). In irradiated cells, the kinetics of mCh-53BP1_ct_ foci accumulation and resolution was strikingly different on soft vs stiff substrates (Fig. 3B, Suppl. Fig. S7E, Suppl. Movies S2-S3). Cells on stiff ECM accumulated more foci and faster compared to cells on soft ECM. Foci resolution was also different. Cells on stiff gels displayed a fast rate of foci resolution during the first six hours post-IR, followed by a slower resolution kinetic, as observed previously [68–70]. In contrast, the rate of foci disappearance was constant for cells in soft growth conditions. Importantly, for cells in stiff conditions, DSB foci returned to pre-IR levels within 12h, whereas cells on soft ECM retained ∼50% of IR-induced foci (Fig. 3B).

While most DSBs have relatively stable positions [71], some break sites coalesce in large repair foci [72–74]. The differences in DSB foci decay curves after IR may therefore reflect an effect of ECM stiffness on DNA repair and/or on DSB clustering. DSB foci fusion events were more frequent in cells growing on stiff (12 kPa) than on soft (0.2 kPa) ECM, with an average of 4.5 vs 1.8 fusion/cell over the 24-hour imaging period, respectively. However, the proportion of DSB foci clustering at any given time point was low (5% or less; Suppl. Fig. S7F) and foci diffusion speed— a likely determinant of clustering—did not differ between stiffness conditions (Suppl. Fig. S7G). Clustering may therefore not fully account for the different foci decay kinetics. Rather, the results indicate that cells repair DSBs faster in a rigid environment, as seen with the comet assay.

### Intermediate filaments define the mechanogenomic response to DNA damage

Comparing individual cells in live imaging data revealed heterogeneity in DSB foci accumulation and resolution, both in soft and stiff contexts, likely reflecting cell-specific characteristics (Suppl. Fig. S7H). In particular, we noticed that a fraction of cells had nuclei with persistent nuclear rotations, ‘spinning’ (Fig. 3C, Suppl. Movie S4), as observed previously in multiple mammalian cell types [75]. Cells with spinning nuclei represented ∼10% and 20% of the population in stiff and soft conditions, respectively (Fig. 3D). Importantly, for these spinning cell nuclei, substrate stiffness did not influence DSB foci accumulation post damage (Fig. 3E). Spinning of the cell nucleus requires an intact cytoskeleton [76], with intermediate filaments (IF) acting as a ‘brake’ for nuclear rotations [77,78]. Hence, the structural continuum coupling the cell nucleus to the ECM [40] may be involved in the response to DNA breaks. We sought to test this hypothesis.

Disruption of the IF network with okadaic acid [79] abolished the effect of ECM stiffness on the DSB response, visualized by 53BP1 foci accumulation (Fig. 4A-B). A similar observation was made using a low concentration of acrylamide, known to disrupt IFs [80] (Fig. 4C-D). In contrast, cells with depolymerized cytoskeletal actin filaments or disrupted microtubules retained a mechano-dependent response to DSBs (Fig. 4A-B). Keratins 8 and 18 are major IF components in luminal breast epithelial cells. Accordingly, co-depletion of K8 and K18 using small hairpin RNAs eliminated the effect of substrate stiffness on 53BP1 foci accumulation following irradiation (Fig. 4E-F). Interestingly, both the acrylamide treatment and K8/18 depletion led to an intermediate level of 53BP1 foci accumulation compared to the soft/stiff controls. Similarly, rotating cells on soft and stiff substrates accumulated mCh-53BP1_ct_ foci at levels corresponding to the intermediate stiffness condition (Fig. 3E). This phenomenon may reflect keratin compensation mechanisms [81] or a dualistic function of IF in the DDR.

**Figure 4.**
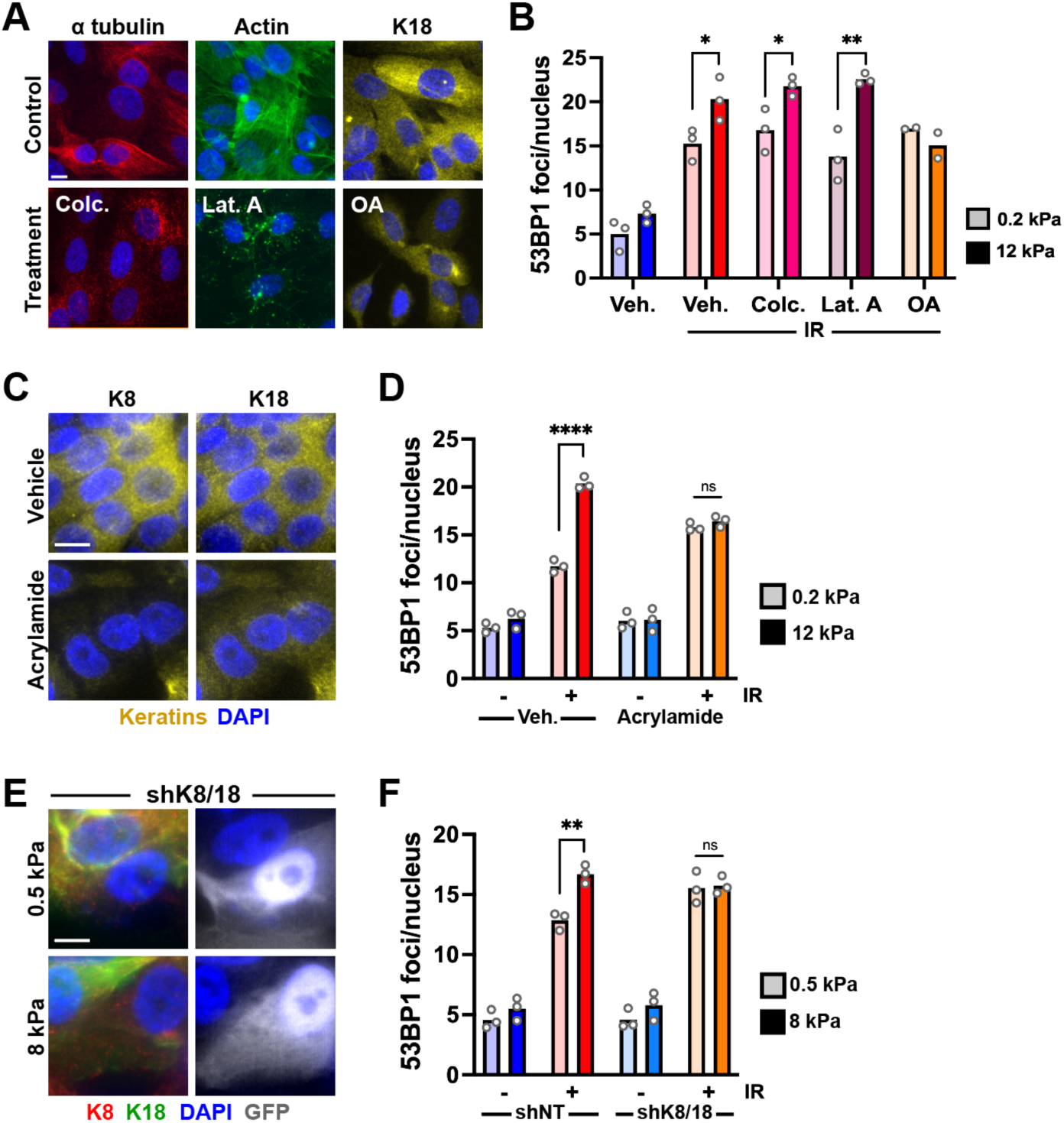
Intermediate filaments modulate the DNA damage response. (**A**) Disruption of cytoskeletal elements in MCF10A cells. Cells were treated with colcemid (0.5 µM; 1h), latrunculine A (1 µM; 90 min), and okadaic acid (OA; 0.1 µM; 2h) to disrupt microtubules, actin filaments, and IFs, respectively. Controls were treated with vehicle. (**B**) IR-induced 53BP1 repair foci in MCF10A cells cultured on soft or stiff PA gels and treated with cytoskeleton-disrupting drugs as in (A). *, *P* < 0.05 and **, *P* < 0.01 (unpaired t-test). (**C**) IF disruption with acrylamide (5 mM; 24h) validated by dual staining of K8 and K18. (**D**) Effect of IF disruption withy acrylamide on IR-induced 53BP1 repair foci accumulation in MCF10A cells on soft or stiff PA substrates. ****, *P* < 0.0001; ns, not significant (unpaired t-test). (**E**) Depletion of K8 and K18 in MCF10A cells. Cells expressing shRNA pools targeting K8 and K18 co-express GFP. (**F**) IR-induced 53BP1 repair foci in cells transduced with nontargeting (NT) shRNA and with shRNA for K8 and K18. Cells were cultured on soft or stiff silicone substrates. Foci scoring was restricted to GFP-positive cells. **, *P* < 0.01; ns, not significant (unpaired t-test). Scale bars, 10 µm.

Next, we examined K8 and K18 expression in different stiffness conditions. Quantitative immunostaining and western blot and analyses of breast epithelial cells confirmed expression of the two keratins in both soft and stiff culture conditions and revealed a shift in relative levels, with an increase in the K8:K18 ratio on stiff substrates (Suppl. Fig. S8A-B). A previous study profiled normal breast tissue transcriptomes using samples from the Komen Tissue Bank [82]. As expected, expression of *KRT8* and *KRT18* was correlated (Suppl. Fig. S8C), and relative transcripts levels (*KRT8*:*KRT18*) scaled with the expression of a 12-gene signature of mammographic density derived from previous studies [48,83,84] (Suppl. Fig. S8D-E). Taken together, the data show that K8/18 intermediate filaments are necessary for the differential DNA damage response of breast epithelial cells in soft and stiff environments and that their relative levels change according to substrate stiffness, which may act as a DDR signaling rheostat.

### ECM adhesion enhances the DDR in stiff environments

Intermediate filaments bind to hemidesmosomal α6/β4 integrins at the plasma membrane—a key determinant of apicobasal polarity in epithelial cells (Fig. 5A). Immunostaining for β4 integrin revealed a strikingly different arrangement of the protein as a function of ECM stiffness, ranging from granular patterns on soft matrices to a quasi-uniform distribution on stiff substrates (Suppl. Fig. S9A). Apicobasal polarity is progressively lost in the HMT-3522 breast cancer progression series [44] (Suppl. Fig. S9B). We therefore compared IR responses in 3D cultures of HMT-3522 S2 (’S2’) cells, a model of ductal carcinoma *in situ*, and of T4-2 cells, forming nodules reminiscent of invasive ductal carcinoma [85]. S2 structures largely retain cell-ECM contacts via basal hemidesmosomal integrins whereas T4-2 nodules are disorganized, with complete loss of apicobasal polarity. Comparing IR-induced accumulation and resolution of 53BP1 DSB foci, we found differences between S2 and T4-2 structures that parallelled the differences between stiff and soft substrates: there was a higher accumulation of DSB foci (30 min time point) and a faster resolution of these foci in S2 compared to T4-2 nodules (Suppl. Fig. S9C). The S2 cell line produces heterogeneous structures [85]. Smaller structures have distinctively basal (’polarized’) hemidesmosomes whereas α6/β4 integrins tends to be mislocalized in larger structures. In polarized S2 nodules, 53BP1 foci accumulation was more pronounced after irradiation than in nonpolarized nodules (Suppl. Fig. S9D).

**Figure 5.**
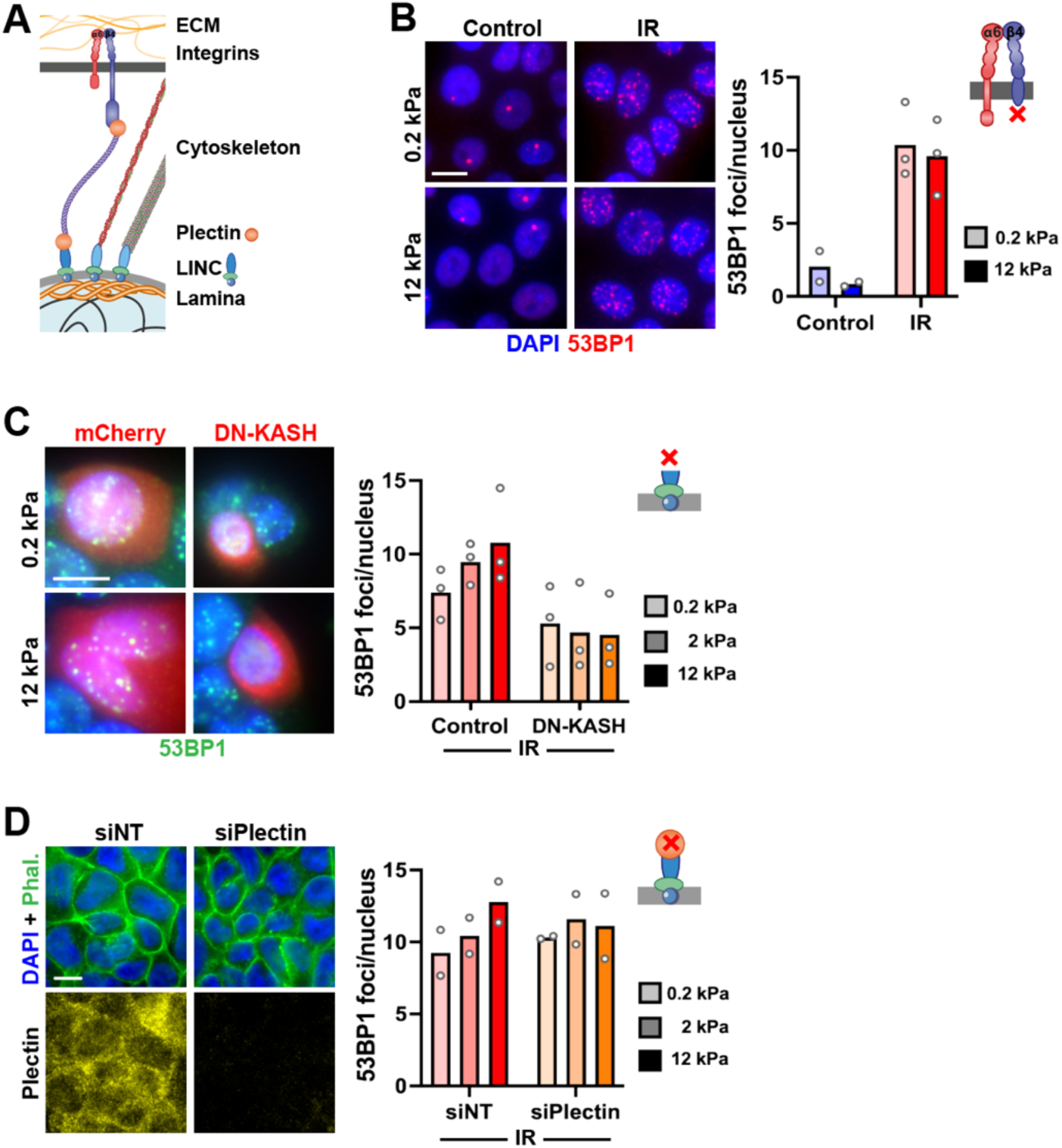
Mechanotransduction regulates the DNA damage response. (**A**) Schematic of the structural continuum between the ECM and the cell nucleus. (**B**) 53BP1 repair foci in S1 β4-’tailless’ cells cultured on soft or stiff PA gels. (**C**) 53BP1 foci post-irradiation in S1 cells transfected with DN-KASH-mCherry or mCherry as control. Foci were scored in mCherry-positive cells. (**D**) 53BP1 foci in irradiated S1 cells transfected with nontargeting siRNA (siNT) or siRNA against plectin. Down-regulation of plectin expression was confirmed by immunostaining. Scale bars, 10 µm.

To further assess the relation between hemidesmosomal integrins and the mechanical aspect of the DNA damage response, we used S1 cells expressing a truncated, cytoplasmic ’tailless’, form of β4 integrin, unable to bind intermediate filaments [45]. In these cells IR-induced accumulation of 53BP1 foci was similar in soft and stiff contexts (Fig. 5B). Collectively, the data highlight a role of hemidesmosomes-IF structural units in genome maintenance.

### The LINC complex is necessary for the mechanoregulation of the DDR

IFs bind the nuclear envelope via the LINC complex which mediates force transmission between the cytoskeleton and the nucleus. Disrupting this interaction with the expression of a dominant negative nesprin KASH mutant (DN-KASH) [86] reduced DSB foci accumulation in irradiated cells and abrogated differences between cells on soft and stiff ECM (Fig. 5C). Similarly, silencing of plectin, which connects IFs to the LINC complex and to integrins, reduced the mechano-dependency for DSB foci accumulation (Fig. 5D). Together, the results indicate that uncoupling IFs from the nuclear envelope disrupts the mechano-dependent DNA damage response.

### Lamins are not altered by ECM stiffness in breast epithelial cells

At the organismal level and in certain cell types (e.g., mesenchymal stem cells), lamin A expression scales with ECM stiffness [87], and laminopathies—genetic disorders driven by *LMNA* mutations—are characterized by genome instability and defective DNA repair [88]. Both A- and B-type lamins interact with DSB factors, including 53BP1. Loss of lamin A/C leads to reduced 53BP1 levels [89,90] and 53BP1 binding to lamin A/C and lamin B1 in the absence of damage has been proposed as a sequestration mechanism, restricting chromatin access at steady state [90,91]. We found no difference in lamin expression levels in breast epithelial cells grown on soft vs. stiff substrates (Suppl. Fig. S10A-B), consistent with a previous report [92]. Lamin localization was similar in the soft and stiff conditions (Suppl. Fig. S10B) and the proportion of lamin A/C relative to lamin B did not vary with ECM stiffness (Suppl. Fig. S10C). We also evaluated publicly available proteomic data from human breast tissue [48] and found constant lamin A/C levels across the four BIRADS breast density categories (Suppl. Fig. S10D). While these findings do not exclude an important role for the lamina as a transducer of mechanical cues to regulate genome maintenance, they indicate that stiffening of the ECM has no dramatic impact on lamin expression and localization in non-neoplastic breast epithelial cells.

### Nuclear YAP is necessary for an efficacious DDR

The YAP/TAZ proteins relay mechanical cues from the ECM to the cell nucleus [93] and ECM stiffness promotes nuclear entry of YAP [94]. Accordingly, S1 breast epithelial cells on a soft substrate had predominantly cytoplasmic YAP whereas the protein localized to the nucleus in cells on stiffer matrices (Fig. 6A). Treatment with the YAP inhibitor verteporfin—which prevents the YAP-TEAD interaction [95] and increases cytoplasmic sequestration by the 14–3-3σ protein [96]— prevented nuclear localization of YAP, even for cells on a stiff substrate (Fig. 6A). In these conditions, the effect of ECM stiffness on DSB repair foci accumulation was lost (Fig. 6B). In MCF10A cells, YAP signals were more heterogeneous. Yet, the proportion of nuclear YAP localization was also greater on stiffer substrates (Suppl. Fig. S11A). Again, verteporfin treatment significantly reduced YAP localization to the nucleus and dampened DSB foci accumulation in irradiated cells (Suppl. Fig. S11B). We conclude that YAP is a mechanosensitive amplifier of the DNA damage response.

**Figure 6.**
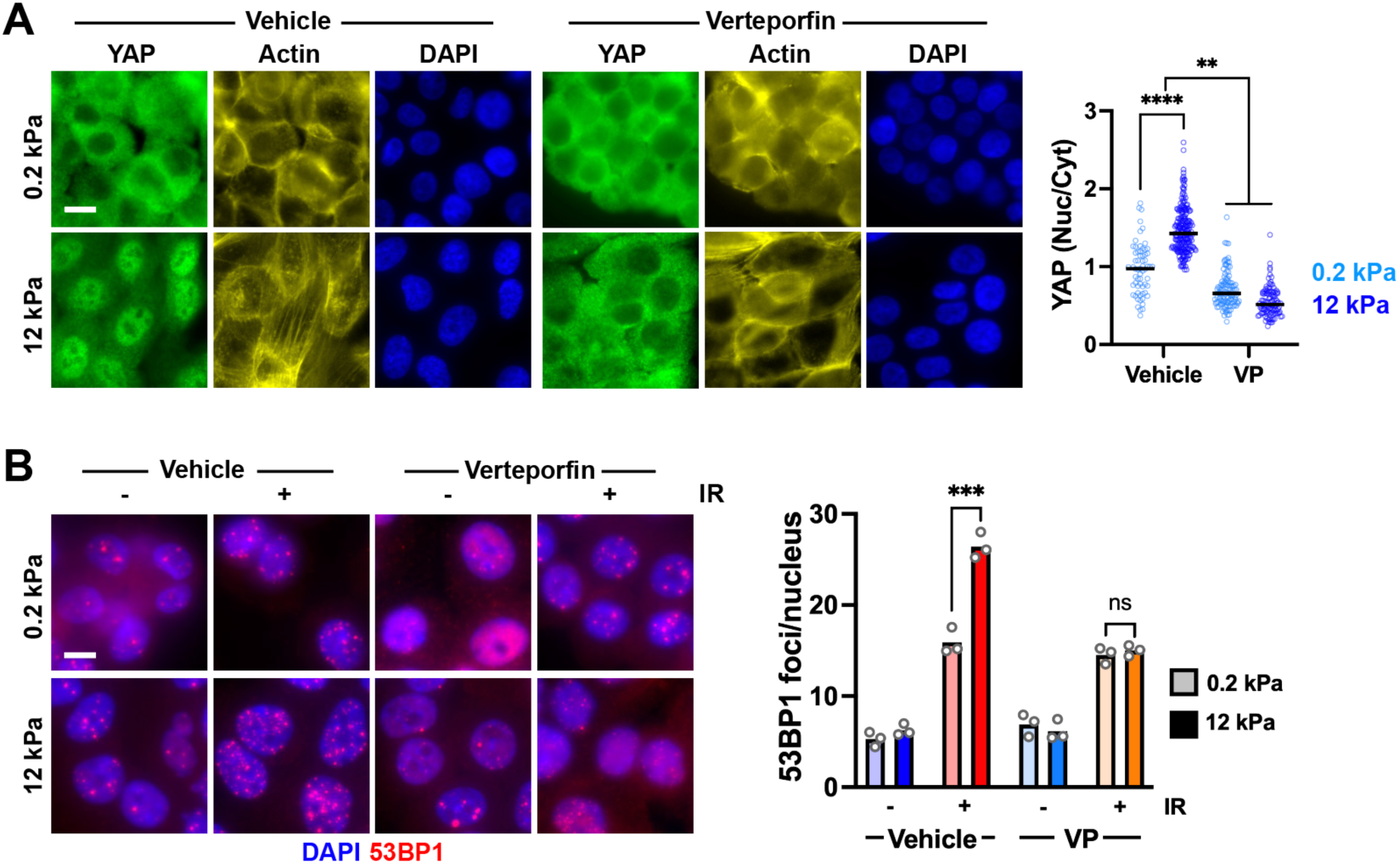
YAP mediates a mechano-dependent DNA damage response. (**A**) YAP localization in S1 cells cultured on soft (0.2 kPa) and stiff (12 kPa) PA gels. Cells were treated with vehicle or verteporfin (VP; 1µM, 6h). Actin and DAPI signals highlight cytoplasmic and nuclear regions, respectively. Quantification of nuclear-to-cytoplasmic YAP intensity ratios is shown in the graph. ****, *P* < 0.0001 and **, *P* < 0.005 (Kruskal-Wallis and Dunn’s multiple comparison test N > 60 cells). (**B**) 53BP1 foci in irradiated and nonirradiated cells treated as in A. ***, *P* = 0.0009; ns, not significant (unpaired t-test).

### Stiff ECM promotes open chromatin landscapes permissive for the DNA damage response

Activated YAP increases chromatin accessibility [97,98] and mechanically-tensed contexts are generally associated with open chromatin states characterized by higher histone acetylation and lower methylation levels [39]. Accordingly, we measured a monotonic increase in the acetylation of histone H3 Lys 9 (H3K9ac) relative tri-methylation at the same site (H3K9me3) as a function of ECM stiffness (Fig. 7A). The coefficient of variation (CV) of DNA signals, reflecting chromatin compaction [57] was also reduced for cells on stiff compared to soft substrates (Fig. 7B). Verteporfin treatment significantly reduced euchromatin levels (H3K9ac) in cells growing on stiff but not on soft substrates (Fig. 7C), linking chromatin relaxation to YAP activity. Importantly, 53BP1 foci accumulation was correlated with H3K9ac levels (Fig. 7D), indicating that ECM stiffness impacts the DDR by altering chromatin organization.

**Figure 7.**
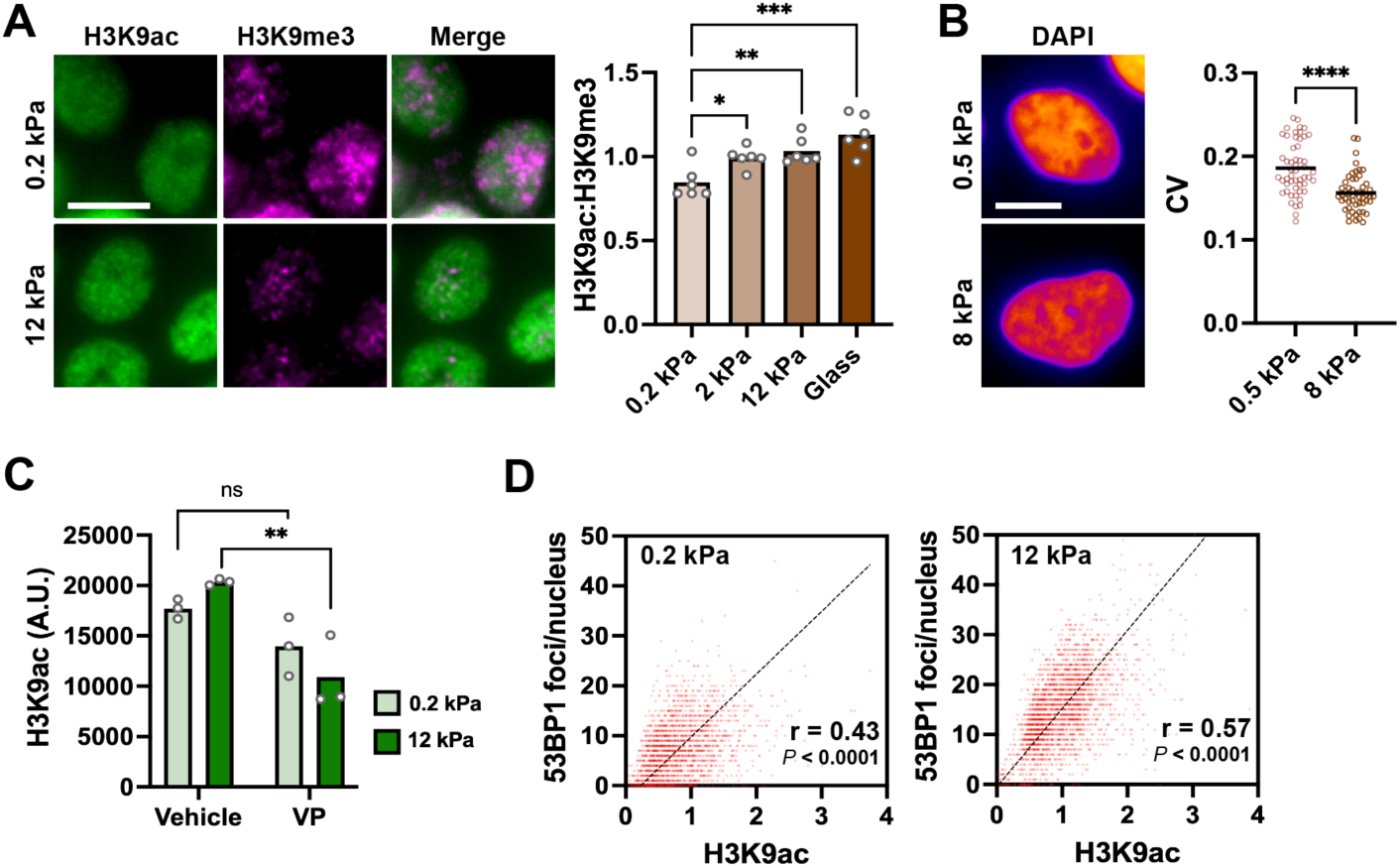
Stiff ECM leads to open chromatin conducive to the DNA damage response. (**A**) Relative levels of H3K9ac and H3K9me3 in S1 cells in various substrate stiffness conditions. ***, *P* = 0.0001; **, *P* < 0.005; *, *P* < 0.05 (ANOVA and Tukey’s multiple comparison test). (**B**) Coefficient of variation (CV) of DAPI signal intensity in MCF10A cells grown on soft (0.5 kPa) and stiffer (8 kPa) silicone substrates. ****, *P* < 0.0001 (Mann-Whitney; N > 50 nuclei). (**C**) H3K9ac expression in S1 cells treated with vehicle or verteporfin (VP; 1µM, 6h). **, *P* < 0.005; ns, not significant (ANOVA and Bonferroni’s multiple comparison test). (**D**) Association between H3K9ac intensity and 53BP1 foci counts in irradiated S1 cells. The Spearman correlation coefficients and associated *P* values are indicated (N > 2000 cells from 3 replicates).

### DNA repair in normal human breast tissue depends on the local mechanical context

We analyzed normal breast tissue to assess the translational relevance of our findings. High-density breast tissue is characterized by high levels of aligned collagen fibrils surrounding mammary ducts [48,84]. As a first approach, collagen alignment was quantified within epithelial regions by picrosirius red staining and birefringence microscopy on normal breast tissue donated to the Komen Tissue Bank. In parallel, DNA damage foci abundance was estimated by 53BP1 immunostaining and proliferation by Ki67 staining (Suppl. Fig. S12A-D). Tissue with the highest collagen alignment tended to have higher levels of 53BP1 foci and higher proliferation, but the differences were not significant. Importantly, intra-tissue heterogeneity was high, as observed previously [9,48]: the standard deviation in collagen alignment between different epithelial regions from a given tissue was comparable to inter-individual variability (Suppl. Fig. S12E). Elastic moduli derived from atomic force microscopy (AFM) measurements of normal breast tissue also highlighted both inter- and intra-individual heterogeneity (Fig. 8A and Suppl. Fig. S12F-H), with modal values of 500-1000 Pa matching values from previous studies [9,48], but marked differences between distinct tissue locations.

**Figure 8.**
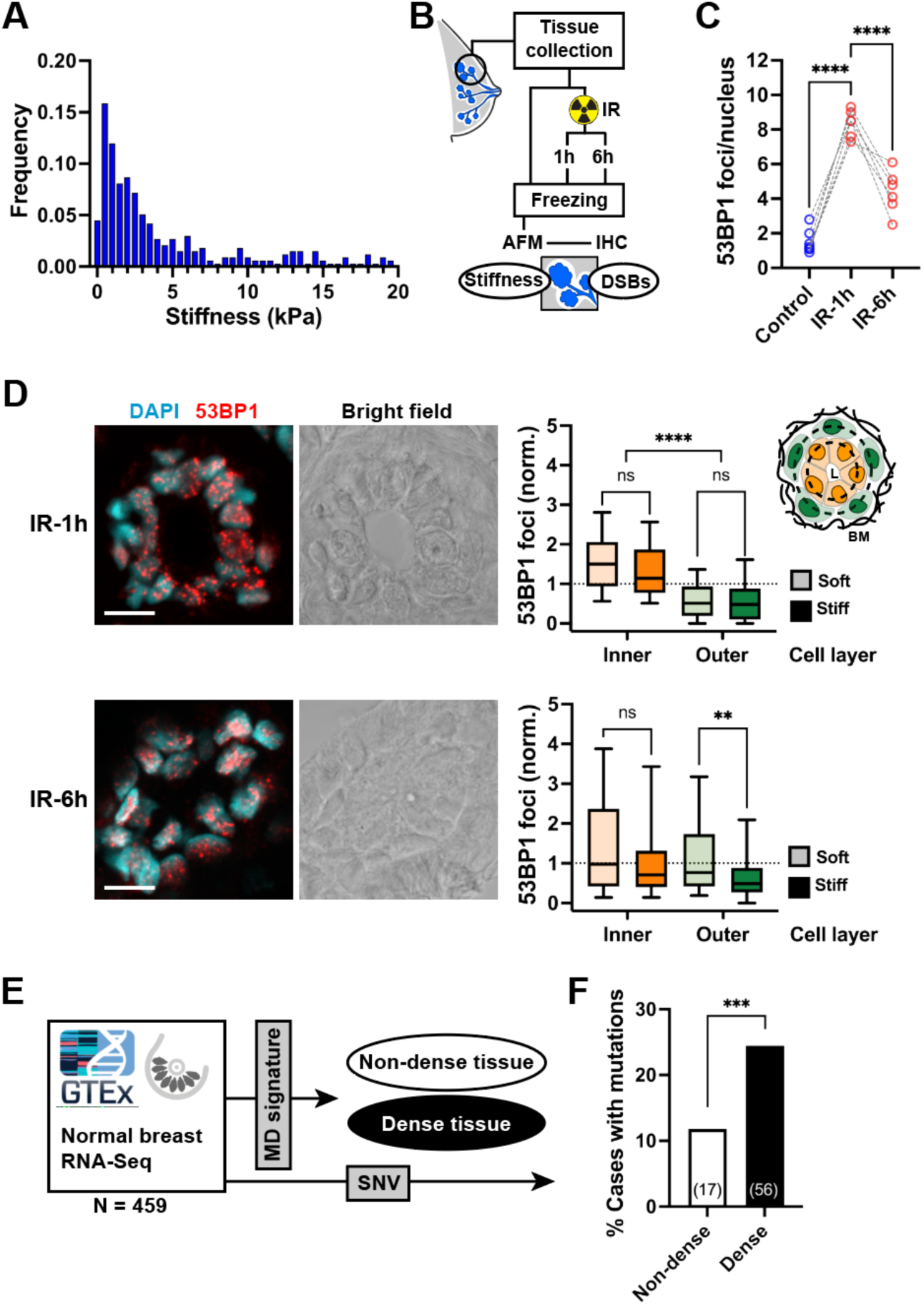
The DNA damage response and mutation load depends on mechanical characteristics of the breast. (**A**) Histogram of breast tissue stiffness values quantified by atomic force microscopy (AFM). N = 334 measurements from 6 breast reduction mammoplasty samples. (**B**) Approach to match AFM measurements around breast epithelial structures and detection of DSB repair foci by immunohistochemistry (IHC) in response to *ex-vivo* irradiation (IR, 3 Gy). (**C**) Average number of IR-induced 53BP1 foci in breast epithelial regions. Controls were mock-irradiated. Dotted lines connect measurements from individual patients. ****, *P* < 0.0001 (ANOVA and Tukey’s multiple comparison test). (**D**) Confocal images of 53BP1 immunostaining (left), and quantification of 53BP1 foci matched to AFM measurements (right). Immunofluorescence images are average intensity projections of 10 optical sections. Scale, 10 µm. Cells were categorized based on their belonging to outer or inner layers, as illustrated in the schematic. Soft and stiff categories were assigned according to median stiffness values. For each time point (1h and 6h post-IR), 53BP1 foci counts were normalized for each patient (n = 6). ****, *P* < 0.0001; **, *P* < 0.0001; ns, not significant (Kruskal-Wallis and Dunn’s multiple comparison test). (**E**) Schematic of the approach. RNA-Seq data from normal breast tissue were used for breast density classification based on median values of a 25-genes pannel. Single nucleotide variations (SNV) were called in the same dataset. (**F**) Proportion of tissues with at least one SNV according the the mammographic density (MD) category. ***, *P* = 0.0006 (Fischer’s exact test).

We therefore devised a strategy to exploit this heterogeneity, relating ECM stiffness to DNA damage induction and repair (Fig. 8B). For this approach, fresh surgical specimens from reduction mammoplasties were cultured short-term *ex vivo*. For each patient, a fraction of tissue was irradiated and left to recover for one or six hours. The epithelium from irradiated tissues had clearly more 53BP1 damage foci than nonirradiated controls, and the number of foci diminished during the recovery period, indicating repair (Fig. 8C). AFM measurements were performed around epithelial structures (lobules and ducts). Then tissue sections were fixed, stained and the same regions were re-analyzed by confocal microscopy, to match the number of DSB foci in cell nuclei to the nearest measurement of tissue stiffness (Suppl. Fig. S12I). The mammary epithelium consists of an inner layer of luminal cells and an outer layer of myoepithelial cells in direct contact with the ECM. IR-induced DSB foci were more abundant 1h post-irradiation in the inner compared to the outer layer (8.1 ± 4.6 vs. 3.0 ± 2.3; averages from all patients). In the inner layer, there was a 22% decrease in DSB foci after recovery, whereas there was no reduction in the global average of repair foci in outer layer. Interestingly, normal breast epithelial cells are less compliant than the ECM. The average stiffness of a mature bilayer of HMEC is 7 kPa [42], whereas the modal elastic modulus of normal breast tissue is 0.6 kPa (Fig. 8A). This difference is visible in AFM maps (Suppl. Fig. S12H). Hence, inner layer cells have their basal side in contact to a stiffer environment than outer layer cells. Inner layer cells also show higher DSB foci accumulation and repair activity than cells from the outer layer, which parallels our observations *in vitro*.

Next, cells were classified into ’soft’ and ’stiff’ categories based on the stiffness of the adjacent ECM (i.e., the closest AFM measurement), using the median as cutoff. We normalized DSB foci counts for each patient to isolate tissue mechanics from other confounders and compared those counts in the stiff and soft groups. There was no difference 1h after irradiation (Fig. 8D, top). At the 6h time point, cell nuclei in a stiffer context had significantly less DSB foci than nuclei in softer environment (Fig. 8D, bottom). This effect was restricted to the outer layer contacting the ECM. These findings indicate that tissue stiffness promote DSB repair.

### Breast tissues with high MD signatures accumulate more mutations

Our data indicate that a stiff cellular microenvironment promotes DNA repair and survival of cells with DNA damage. As a corollary to this pro-survival effect, cells in rigid conditions may accumulate mutations. To test this hypothesis, we retrieved RNA-Seq data from normal breast tissue (N = 459). Tissues were classified as ’low’ and ’high’ mammographic density according to a gene signature established previously [59] (Fig. 8E). Age and BMI were not different in the high versus low MD groups (Suppl. Fig. S13A). Reactome pathway analysis [99] identified lipid metabolism processes among low-expressed genes from the high MD group (Suppl. Table S1), which was expected given that high-density breasts tend to have lower adiposity. Pathways related to cell junctions were enriched when considering over-expressed genes from the high MD group (Suppl. Table S2). Mutations were called in ∼20% of the tissues. These were mostly miss-sense mutations, with a bias toward C>T single nucleotide variations (Suppl. Fig. S13B-C). Importantly, the frequency of mutations was more than two-fold higher in high-MD compared to low-MD tissues (Fig. 8F).

## Discussion

Tissue mechanics impacts genome integrity. Cell migration in confined spaces compresses and bends cell nuclei to the point of nuclear envelope rupture [40], leading to an exodus of repair factors and nuclear entry of cytoplasmic endonucleases causing acute DNA damage [100–102]. Nuclear deformations also cause replication stress and DNA breaks, even without nuclear rupture [103]. While metastasizing cancer cell nuclei may be subjected to these extreme distortions, most normal epithelia experience more subtle mechanical stimuli. Here, we show that stiff cell environments promote repair of DNA breaks via direct force transmission across the structural continuum spanning integrins, intermediate filaments, and the LINC complex. This effect is mediated by YAP which promotes an open chromatin organization permissive for the DNA damage response.

We find that enhanced DSB response in stiff conditions is a general effect, observed in noncancerous and cancerous epithelial cells from different origins. The scaling relationship between the elastic modulus of the substrate and the DDR seems, however, to be context specific. The effect was monotonic for non-neoplastic breast epithelial cells, plateaued at 2 kPa in breast cancer cells, was shifted towards higher moduli in hepatic cells, and was not detected in osteosarcoma cells, which likely reflects mechanical tuning to distinct microenvironments.

Cell-ECM contacts via integrins are determinant for DNA repair and cell survival [34,104–107]. The basal distribution of hemidesmosomal integrins was strongly influenced by substrate stiffness (as seen previously [11]), and expression of a tailless β4 integrin mutant supressed mechano-dependency for the DDR. In epithelial cells, α6/β4 integrins are particular in that they bind the IF network rather than actin filaments. We show that pharmacological disruption of IFs or depletion of major components of the IF network in breast luminal cells eliminate the DDR mechano-dependency. Stiff ECM as well as compressive strain have been shown to result in more interconnected and more rigid IF networks that may directly impact nuclear mechanosignalling [36,37]. Although we do not exclude a direct contribution of IF proteins in the DDR in breast epithelial cells (as shown for keratin 17 [108]), K8/18 have not been localized to the cell nucleus [109]. A DDR rheometer mechanism involving the cytoplasmic keratin network is therefore more likely. Keratin filaments are obligate heteropolymers and K8/K18 is the canonical pair in sinple epithelia [110]. We found that the *KRT8*:*KRT18* expression ratio increases with substrate stiffness in breast epithelial cell lines and scales with markers of MD in normal breast tissue. The impact of an excess of K8 on the organization and mechanical properties of the network deserves further investigation.

Altered force transmission to the nucleus (as a consequence of the remodeling of hemidesmosome-keratin structural units) impacts on YAP signaling [94,111]. As expected, YAP was predominantly nuclear in cells on stiffer substrates and cytoplasmic in cells on softer matrices. Excluding YAP from the nucleus reduced the DDR, highlighting YAP as a mechanosensitive modulator of the DDR. This finding contrasts with a previous study by Deng et al. [112] who found an effect of ECM stiffness on the DDR independent of YAP. The role of YAP in genome maintenance is therefore likely context-dependent [113–115]. Expression of oncogenic YAP (i.e., constitutively active, nuclear YAP) leads to broad changes in chromatin accessibility and results in a net opening of chromatin in breast epithelial cells [98]. Accordingly, cells on stiffer substrates had globally more opened chromatin. Verteporfin reduced euchromatin marks and blunted the DDR.

DNA accessibility is a key determinant of repair efficacy and outcomes, with slower DSB repair in heterochromatin than in euchromatin [116–119]. Stiffness has an impact on chromatin organization and cells in stiff ECM tend to have more open chromatin [120–122] (Fig. 7) that may explain, at least in part, their faster repair kinetics. The DDR is indeed amplified in the context of euchromatin [123,124]. Our study identifies an effect of ECM stiffness on early DDR events, i.e. the stabilization of ATM at DSBs. Chromatin relaxation promotes the formation of R-loops contributing to ATM activation [125,126]. It will hence be interesting to assess mechanogenomic effects on secondary DNA structures tied to the DDR.

This study has limitations. Human breast tissues were derived from reduction mammoplasties which have similarities with benign breast disease [127] and can therefore not be considered fully normal. This may in part explain why tissue elastic moduli (median = 2.7 kPa) were higher than in some [9,11] (but not all [8]) previous studies. Also, we could not control for potential confounding factors such as obesity which induces fibrosis [14,128] as well as elevated levels of DSBs in the breast epithelium [129]. We therefore normalized DNA damage values for each patient, exploiting intra-individual heterogeneity in tissue mechanics to highlight local effects of tissue stiffness on the DNA damage response. We will consider potential synergistic effects of tissue density, obesity, and genetic cancer predisposition on genome integrity in future studies.

ECM stiffness is a multifactorial driver for malignant phenotypes. Stiff microenvironments alter gene expression, re-wire biochemical signaling, and promote the expansion of breast epithelial stem/progenitor cells [130–133]. Genomic instability in stiff breast tissue may also contribute to elevated cancer risk conferred by high MD. Previous studies have shown that increased ECM stiffness promotes DNA damage in mammary epithelial cells through at least two mechanisms: the accumulation of reactive aldehydes [22,134] and altered mitochondrial function leading to elevated ROS levels [135]. Both are consistent with the increased levels of DNA breaks observed in high MD tissue [22,136] and in hepatocytes in stiff ECM [137], as well as elevated levels of point mutations in tissues with high MD signatures (Fig. 8F) and in stiff breast tumors [22]. Our study highlights a new mechanism linking DNA damage to tissue rigidity in which stiff ECM exacerbates the DNA damage response and promotes DNA repair. We stress that, in breast acini, significant differences in the DDR were measured for a relatively modest (60%) increase in ECM stiffness. This increase parallels the 1.5 to 2-fold augmentation of the Young’s modulus in high- vs. low-MD tissue [48]. This finding may seem counterintuitive, contradicting a general consensus that *lower* DNA repair capability increases (breast) cancer risk [138]. Yet, mutagenic DSB repair is associated with breast cancer initiation [20]. In a stiff context with heightened DNA damage, cells repairing their DNA damage at the expense of apoptosis will accumulate mutations–as seen in high MD normal breast tissue in this study–favoring pre-cancerous genome instability in a pro-oncogenic microenvironment.

## Data availability

All data will be made available upon request once published.

## Supplementary data

Supplementary data is available at NAR Online.

## Acknowledgments

We thank Hamza Rashid, Tucker Loy, and Matt Gray for technical assistance, Thierry Pécot for help with image analyses, Dr. Kurt Hodges and Dr. Ayaz Shaikh for their active involvement in the breast tissue collection, Dr. Valerie Weaver and Dr. Mina Bissell for cell lines, Dr. Daniel Conway for the DN-KASH construct, and Dr. Titia de Lange for the mCh-53BP1_ct_ construct. The data used for RNAseq analyses were retrieved from the portal of the GTEx Project.

## Author contributions

conceptualization: PAV, SL; investigation: CD, ML, CL, AS, HL, JL, QS, MG, SS, MB; supervision: PAV, KB, MG; writing—original draft: PAV; writing—review & editing: CD, ML, SL, KB, MG.

## References

1. Byrne, C., Schairer, C., Wolfe, J., Parekh, N., Salane, M., Brinton, L.A., Hoover, R. and Haile, R. (1995) Mammographic features and breast cancer risk: effects with time, age, and menopause status. J Natl Cancer Inst, 87, 1622–1629.

2. McCormack, V.A. and dos Santos Silva, I. (2006) Breast density and parenchymal patterns as markers of breast cancer risk: a meta-analysis. Cancer Epidemiol Biomarkers Prev, 15, 1159–1169.

3. Pettersson, A., Graff, R.E., Ursin, G., Santos Silva, I.D., McCormack, V., Baglietto, L., Vachon, C., Bakker, M.F., Giles, G.G., Chia, K.S. et al. (2014) Mammographic density phenotypes and risk of breast cancer: a meta-analysis. J Natl Cancer Inst, 106.

4. Bodewes, F.T.H., van Asselt, A.A., Dorrius, M.D., Greuter, M.J.W. and de Bock, G.H. (2022) Mammographic breast density and the risk of breast cancer: A systematic review and meta-analysis. Breast, 66, 62–68.

5. Bertrand, K.A., Tamimi, R.M., Scott, C.G., Jensen, M.R., Pankratz, V., Visscher, D., Norman, A., Couch, F., Shepherd, J., Fan, B. et al. (2013) Mammographic density and risk of breast cancer by age and tumor characteristics. Breast Cancer Res, 15, R104.

6. Yaghjyan, L., Pettersson, A., Colditz, G.A., Collins, L.C., Schnitt, S.J., Beck, A.H., Rosner, B., Vachon, C. and Tamimi, R.M. (2015) Postmenopausal mammographic breast density and subsequent breast cancer risk according to selected tissue markers. Br J Cancer, 113, 1104–1113.

7. Li, T., Sun, L., Miller, N., Nicklee, T., Woo, J., Hulse-Smith, L., Tsao, M.S., Khokha, R., Martin, L. and Boyd, N. (2005) The association of measured breast tissue characteristics with mammographic density and other risk factors for breast cancer. Cancer Epidemiol Biomarkers Prev, 14, 343–349.

8. Samani, A., Zubovits, J. and Plewes, D. (2007) Elastic moduli of normal and pathological human breast tissues: an inversion-technique-based investigation of 169 samples. Phys Med Biol, 52, 1565–1576.

9. Plodinec, M., Loparic, M., Monnier, C.A., Obermann, E.C., Zanetti-Dallenbach, R., Oertle, P., Hyotyla, J.T., Aebi, U., Bentires-Alj, M., Lim, R.Y. et al. (2012) The nanomechanical signature of breast cancer. Nat Nanotechnol, 7, 757–765.

10. Boyd, N.F., Li, Q., Melnichouk, O., Huszti, E., Martin, L.J., Gunasekara, A., Mawdsley, G., Yaffe, M.J. and Minkin, S. (2014) Evidence that breast tissue stiffness is associated with risk of breast cancer. PLoS One, 9, e100937.

11. Paszek, M.J., Zahir, N., Johnson, K.R., Lakins, J.N., Rozenberg, G.I., Gefen, A., Reinhart-King, C.A., Margulies, S.S., Dembo, M., Boettiger, D. et al. (2005) Tensional homeostasis and the malignant phenotype. Cancer Cell, 8, 241–254.

12. Levental, K.R., Yu, H., Kass, L., Lakins, J.N., Egeblad, M., Erler, J.T., Fong, S.F., Csiszar, K., Giaccia, A., Weninger, W. et al. (2009) Matrix crosslinking forces tumor progression by enhancing integrin signaling. Cell, 139, 891–906.

13. Wei, S.C., Fattet, L., Tsai, J.H., Guo, Y., Pai, V.H., Majeski, H.E., Chen, A.C., Sah, R.L., Taylor, S.S., Engler, A.J. et al. (2015) Matrix stiffness drives epithelial-mesenchymal transition and tumour metastasis through a TWIST1-G3BP2 mechanotransduction pathway. Nat Cell Biol, 17, 678–688.

14. Seo, B.R., Bhardwaj, P., Choi, S., Gonzalez, J., Andresen Eguiluz, R.C., Wang, K., Mohanan, S., Morris, P.G., Du, B., Zhou, X.K. et al. (2015) Obesity-dependent changes in interstitial ECM mechanics promote breast tumorigenesis. Sci Transl Med, 7, 301ra130.

15. Carey, S.P., Martin, K.E. and Reinhart-King, C.A. (2017) Three-dimensional collagen matrix induces a mechanosensitive invasive epithelial phenotype. Sci Rep, 7, 42088.

16. Finger, A.M., Hendley, A.M., Figueroa, D., Gonzalez, H. and Weaver, V.M. (2025) Tissue mechanics in tumor heterogeneity and aggression. Trends Cancer.

17. Ciccia, A. and Elledge, S.J. (2010) The DNA damage response: making it safe to play with knives. Mol Cell, 40, 179–204.

18. Smith, T.R., Miller, M.S., Lohman, K.K., Case, L.D. and Hu, J.J. (2003) DNA damage and breast cancer risk. Carcinogenesis, 24, 883–889.

19. Kennedy, D.O., Agrawal, M., Shen, J., Terry, M.B., Zhang, F.F., Senie, R.T., Motykiewicz, G. and Santella, R.M. (2005) DNA repair capacity of lymphoblastoid cell lines from sisters discordant for breast cancer. J Natl Cancer Inst, 97, 127–132.

20. Keimling, M., Deniz, M., Varga, D., Stahl, A., Schrezenmeier, H., Kreienberg, R., Hoffmann, I., Konig, J. and Wiesmuller, L. (2012) The power of DNA double-strand break (DSB) repair testing to predict breast cancer susceptibility. FASEB J, 26, 2094–2104.

21. DeFilippis, R.A., Fordyce, C., Patten, K., Chang, H., Zhao, J., Fontenay, G.V., Kerlikowske, K., Parvin, B. and Tlsty, T.D. (2014) Stress signaling from human mammary epithelial cells contributes to phenotypes of mammographic density. Cancer Res, 74, 5032–5044.

22. Hayward, M.K., Northey, J.J., Opazo-Mellado, V., Stashko, C., Maller, O., Ironside, A.J., Que, X., Lakins, J.N., Hwang, E.S., Witztum, J.L. et al. (2026) Tissue tension fosters macrophage-driven lipid peroxidation-induced DNA damage. Cancer Cell.

23. Bosch-Fortea, M. and Martin-Belmonte, F. (2018) Mechanosensitive adhesion complexes in epithelial architecture and cancer onset. Curr Opin Cell Biol, 50, 42–49.

24. Kechagia, J.Z., Ivaska, J. and Roca-Cusachs, P. (2019) Integrins as biomechanical sensors of the microenvironment. Nat Rev Mol Cell Biol, 20, 457–473.

25. Simon, D.N. and Wilson, K.L. (2011) The nucleoskeleton as a genome-associated dynamic ’network of networks’. Nat Rev Mol Cell Biol, 12, 695–708.

26. Maniotis, A.J., Chen, C.S. and Ingber, D.E. (1997) Demonstration of mechanical connections between integrins, cytoskeletal filaments, and nucleoplasm that stabilize nuclear structure. Proc Natl Acad Sci U S A, 94, 849–854.

27. Streuli, C.H. (2016) Integrins as architects of cell behavior. Mol Biol Cell, 27, 2885–2888.

28. Barcellos-Hoff, M.H., Aggeler, J., Ram, T.G. and Bissell, M.J. (1989) Functional differentiation and alveolar morphogenesis of primary mammary cultures on reconstituted basement membrane. Development, 105, 223–235.

29. Boudreau, N., Sympson, C.J., Werb, Z. and Bissell, M.J. (1995) Suppression of ICE and apoptosis in mammary epithelial cells by extracellular matrix. Science, 267, 891–893.

30. Weaver, V.M., Lelievre, S., Lakins, J.N., Chrenek, M.A., Jones, J.C., Giancotti, F., Werb, Z. and Bissell, M.J. (2002) beta4 integrin-dependent formation of polarized three-dimensional architecture confers resistance to apoptosis in normal and malignant mammary epithelium. Cancer Cell, 2, 205–216.

31. Zahir, N., Lakins, J.N., Russell, A., Ming, W., Chatterjee, C., Rozenberg, G.I., Marinkovich, M.P. and Weaver, V.M. (2003) Autocrine laminin-5 ligates alpha6beta4 integrin and activates RAC and NFkappaB to mediate anchorage-independent survival of mammary tumors. J Cell Biol, 163, 1397–1407.

32. Sternlicht, M.D., Lochter, A., Sympson, C.J., Huey, B., Rougier, J.P., Gray, J.W., Pinkel, D., Bissell, M.J. and Werb, Z. (1999) The stromal proteinase MMP3/stromelysin-1 promotes mammary carcinogenesis. Cell, 98, 137–146.

33. Radisky, D.C., Levy, D.D., Littlepage, L.E., Liu, H., Nelson, C.M., Fata, J.E., Leake, D., Godden, E.L., Albertson, D.G., Nieto, M.A. et al. (2005) Rac1b and reactive oxygen species mediate MMP-3-induced EMT and genomic instability. Nature, 436, 123–127.

34. Vidi, P.A., Chandramouly, G., Gray, M., Wang, L., Liu, E., Kim, J.J., Roukos, V., Bissell, M.J., Moghe, P.V. and Lelievre, S.A. (2012) Interconnected contribution of tissue morphogenesis and the nuclear protein NuMA to the DNA damage response. J Cell Sci, 125, 350–361.

35. Seltmann, K., Fritsch, A.W., Kas, J.A. and Magin, T.M. (2013) Keratins significantly contribute to cell stiffness and impact invasive behavior. Proc Natl Acad Sci U S A, 110, 18507–18512.

36. Laly, A.C., Sliogeryte, K., Pundel, O.J., Ross, R., Keeling, M.C., Avisetti, D., Waseem, A., Gavara, N. and Connelly, J.T. (2021) The keratin network of intermediate filaments regulates keratinocyte rigidity sensing and nuclear mechanotransduction. Sci Adv, 7.

37. Kechagia, Z., Saez, P., Gomez-Gonzalez, M., Canales, B., Viswanadha, S., Zamarbide, M., Andreu, I., Koorman, T., Beedle, A.E.M., Elosegui-Artola, A. et al. (2023) The laminin-keratin link shields the nucleus from mechanical deformation and signalling. Nat Mater, 22, 1409–1420.

38. Engler, A.J., Sen, S., Sweeney, H.L. and Discher, D.E. (2006) Matrix elasticity directs stem cell lineage specification. Cell, 126, 677–689.

39. Wagh, K., Ishikawa, M., Garcia, D.A., Stavreva, D.A., Upadhyaya, A. and Hager, G.L. (2021) Mechanical Regulation of Transcription: Recent Advances. Trends Cell Biol, 31, 457–472.

40. Kalukula, Y., Stephens, A.D., Lammerding, J. and Gabriele, S. (2022) Mechanics and functional consequences of nuclear deformations. Nat Rev Mol Cell Biol, 23, 583–602.

41. Shivashankar, G.V. (2023) Mechanical forces and the 3D genome. Curr Opin Struct Biol, 83, 102728.

42. Lee, H., Bonin, K. and Guthold, M. (2021) Human mammary epithelial cells in a mature, stratified epithelial layer flatten and stiffen compared to single and confluent cells. Biochim Biophys Acta Gen Subj, 1865, 129891.

43. Bredfeldt, J.S., Liu, Y., Conklin, M.W., Keely, P.J., Mackie, T.R. and Eliceiri, K.W. (2014) Automated quantification of aligned collagen for human breast carcinoma prognosis. J Pathol Inform, 5, 28.

44. Vidi, P.A., Bissell, M.J. and Lelievre, S.A. (2013) Three-dimensional culture of human breast epithelial cells: the how and the why. Methods Mol Biol, 945, 193–219.

45. Spinardi, L., Einheber, S., Cullen, T., Milner, T.A. and Giancotti, F.G. (1995) A recombinant tail-less integrin beta 4 subunit disrupts hemidesmosomes, but does not suppress alpha 6 beta 4-mediated cell adhesion to laminins. J Cell Biol, 129, 473–487.

46. Dimitrova, N., Chen, Y.C., Spector, D.L. and de Lange, T. (2008) 53BP1 promotes non-homologous end joining of telomeres by increasing chromatin mobility. Nature, 456, 524–528.

47. Locatelli, M., Lawrimore, J., Lin, H., Sanaullah, S., Seitz, C., Segall, D., Kefer, P., Salvador Moreno, N., Lietz, B., Anderson, R. et al. (2022) DNA damage reduces heterogeneity and coherence of chromatin motions. Proc Natl Acad Sci U S A, 119, e2205166119.

48. Northey, J.J., Barrett, A.S., Acerbi, I., Hayward, M.K., Talamantes, S., Dean, I.S., Mouw, J.K., Ponik, S.M., Lakins, J.N., Huang, P.J. et al. (2020) Stiff stroma increases breast cancer risk by inducing the oncogene ZNF217. J Clin Invest.

49. Yeung, T., Georges, P.C., Flanagan, L.A., Marg, B., Ortiz, M., Funaki, M., Zahir, N., Ming, W., Weaver, V. and Janmey, P.A. (2005) Effects of substrate stiffness on cell morphology, cytoskeletal structure, and adhesion. Cell Motil Cytoskeleton, 60, 24–34.

50. Sader, J.E., Chon, J.W.M. and Mulvaney, P. (1999) Calibration of rectangular atomic force microscope cantilevers. Review of Scientific Instruments, 70, 3967–3969.

51. Takigawa, T., Morino, Y., Urayama, K. and Masuda, T. (1996) Poisson’s ratio of polyacrylamide (PAAm) gels. Polymer Gels and Networks, 4, 1–5.

52. Prajapati, S., Locatelli, M., Sawyer, C., Holmes, J., Bonin, K., Black, P. and Vidi, P.A. (2022) Characterization and implementation of a miniature X-ray system for live cell microscopy. Mutat Res, 824, 111772.

53. Gyori, B.M., Venkatachalam, G., Thiagarajan, P.S., Hsu, D. and Clement, M.V. (2014) OpenComet: an automated tool for comet assay image analysis. Redox Biol, 2, 457–465.

54. Stringer, C., Wang, T., Michaelos, M. and Pachitariu, M. (2021) Cellpose: a generalist algorithm for cellular segmentation. Nat Methods, 18, 100–106.

55. Roukos, V., Pegoraro, G., Voss, T.C. and Misteli, T. (2015) Cell cycle staging of individual cells by fluorescence microscopy. Nat Protoc, 10, 334–348.

56. Schmidt, U., Weigert, M., Broaddus, C. and Myers, G. (2018). Springer International Publishing, Cham, pp. 265–273.

57. Neguembor, M.V., Martin, L., Castells-Garcia, A., Gomez-Garcia, P.A., Vicario, C., Carnevali, D., AlHaj Abed, J., Granados, A., Sebastian-Perez, R., Sottile, F. et al. (2021) Transcription-mediated supercoiling regulates genome folding and loop formation. Mol Cell, 81, 3065–3081 e3012.

58. Consortium, G.T. (2013) The Genotype-Tissue Expression (GTEx) project. Nat Genet, 45, 580–585.

59. Yang, W.T., Lewis, M.T., Hess, K., Wong, H., Tsimelzon, A., Karadag, N., Cairo, M., Wei, C., Meric-Bernstam, F., Brown, P. et al. (2010) Decreased TGFbeta signaling and increased COX2 expression in high risk women with increased mammographic breast density. Breast Cancer Res Treat, 119, 305–314.

60. Robinson, M.D., McCarthy, D.J. and Smyth, G.K. (2010) edgeR: a Bioconductor package for differential expression analysis of digital gene expression data. Bioinformatics, 26, 139–140.

61. Yu, G., Wang, L.G., Han, Y. and He, Q.Y. (2012) clusterProfiler: an R package for comparing biological themes among gene clusters. OMICS, 16, 284–287.

62. Grafton, M.M., Wang, L., Vidi, P.A., Leary, J. and Lelievre, S.A. (2011) Breast on-a-chip: mimicry of the channeling system of the breast for development of theranostics. Integr Biol (Camb), 3, 451–459.

63. Fischer, R.S., Myers, K.A., Gardel, M.L. and Waterman, C.M. (2012) Stiffness-controlled three-dimensional extracellular matrices for high-resolution imaging of cell behavior. Nat Protoc, 7, 2056–2066.

64. Lopez, J.I., Kang, I., You, W.K., McDonald, D.M. and Weaver, V.M. (2011) In situ force mapping of mammary gland transformation. Integr Biol (Camb), 3, 910–921.

65. So, S., Davis, A.J. and Chen, D.J. (2009) Autophosphorylation at serine 1981 stabilizes ATM at DNA damage sites. J Cell Biol, 187, 977–990.

66. Zgheib, O., Pataky, K., Brugger, J. and Halazonetis, T.D. (2009) An oligomerized 53BP1 tudor domain suffices for recognition of DNA double-strand breaks. Mol Cell Biol, 29, 1050–1058.

67. Fradet-Turcotte, A., Canny, M.D., Escribano-Diaz, C., Orthwein, A., Leung, C.C., Huang, H., Landry, M.C., Kitevski-LeBlanc, J., Noordermeer, S.M., Sicheri, F. et al. (2013) 53BP1 is a reader of the DNA-damage-induced H2A Lys 15 ubiquitin mark. Nature, 499, 50–54.

68. Olive, P.L. and Banath, J.P. (2004) Phosphorylation of histone H2AX as a measure of radiosensitivity. Int J Radiat Oncol Biol Phys, 58, 331–335.

69. Riballo, E., Kuhne, M., Rief, N., Doherty, A., Smith, G.C., Recio, M.J., Reis, C., Dahm, K., Fricke, A., Krempler, A. et al. (2004) A pathway of double-strand break rejoining dependent upon ATM, Artemis, and proteins locating to gamma-H2AX foci. Mol Cell, 16, 715–724.

70. Shibata, A., Conrad, S., Birraux, J., Geuting, V., Barton, O., Ismail, A., Kakarougkas, A., Meek, K., Taucher-Scholz, G., Lobrich, M. et al. (2011) Factors determining DNA double-strand break repair pathway choice in G2 phase. EMBO J, 30, 1079–1092.

71. Soutoglou, E., Dorn, J.F., Sengupta, K., Jasin, M., Nussenzweig, A., Ried, T., Danuser, G. and Misteli, T. (2007) Positional stability of single double-strand breaks in mammalian cells. Nat Cell Biol, 9, 675–682.

72. Aymard, F., Aguirrebengoa, M., Guillou, E., Javierre, B.M., Bugler, B., Arnould, C., Rocher, V., Iacovoni, J.S., Biernacka, A., Skrzypczak, M. et al. (2017) Genome-wide mapping of long-range contacts unveils clustering of DNA double-strand breaks at damaged active genes. Nat Struct Mol Biol, 24, 353–361.

73. Kilic, S., Lezaja, A., Gatti, M., Bianco, E., Michelena, J., Imhof, R. and Altmeyer, M. (2019) Phase separation of 53BP1 determines liquid-like behavior of DNA repair compartments. EMBO J, 38, e101379.

74. Chiolo, I., Altmeyer, M., Legube, G. and Mekhail, K. (2025) Nuclear and genome dynamics underlying DNA double-strand break repair. Nat Rev Mol Cell Biol.

75. Maninova, M., Iwanicki, M.P. and Vomastek, T. (2014) Emerging role for nuclear rotation and orientation in cell migration. Cell Adh Migr, 8, 42–48.

76. Ji, J.Y., Lee, R.T., Vergnes, L., Fong, L.G., Stewart, C.L., Reue, K., Young, S.G., Zhang, Q., Shanahan, C.M. and Lammerding, J. (2007) Cell nuclei spin in the absence of lamin b1. J Biol Chem, 282, 20015–20026.

77. Hay, M. and De Boni, U. (1991) Chromatin motion in neuronal interphase nuclei: changes induced by disruption of intermediate filaments. Cell Motil Cytoskeleton, 18, 63–75.

78. Gerashchenko, M.V., Chernoivanenko, I.S., Moldaver, M.V. and Minin, A.A. (2009) Dynein is a motor for nuclear rotation while vimentin IFs is a “brake”. Cell Biol Int, 33, 1057–1064.

79. Strnad, P., Windoffer, R. and Leube, R.E. (2001) In vivo detection of cytokeratin filament network breakdown in cells treated with the phosphatase inhibitor okadaic acid. Cell Tissue Res, 306, 277–293.

80. Eckert, B.S. (1985) Alteration of intermediate filament distribution in PtK1 cells by acrylamide. Eur J Cell Biol, 37, 169–174.

81. Kanaji, N., Bandoh, S., Fujita, J., Ishii, T., Ishida, T. and Kubo, A. (2007) Compensation of type I and type II cytokeratin pools in lung cancer. Lung Cancer, 55, 295–302.

82. Kang, T., Yau, C., Wong, C.K., Sanborn, J.Z., Newton, Y., Vaske, C., Benz, S.C., Krings, G., Camarda, R., Henry, J.E. et al. (2020) A risk-associated Active transcriptome phenotype expressed by histologically normal human breast tissue and linked to a pro-tumorigenic adipocyte population. Breast Cancer Res, 22, 81.

83. Alowami, S., Troup, S., Al-Haddad, S., Kirkpatrick, I. and Watson, P.H. (2003) Mammographic density is related to stroma and stromal proteoglycan expression. Breast Cancer Res, 5, R129–135.

84. McConnell, J.C., O’Connell, O.V., Brennan, K., Weiping, L., Howe, M., Joseph, L., Knight, D., O’Cualain, R., Lim, Y., Leek, A. et al. (2016) Increased peri-ductal collagen micro-organization may contribute to raised mammographic density. Breast Cancer Res, 18, 5.

85. Rizki, A., Weaver, V.M., Lee, S.Y., Rozenberg, G.I., Chin, K., Myers, C.A., Bascom, J.L., Mott, J.D., Semeiks, J.R., Grate, L.R. et al. (2008) A human breast cell model of preinvasive to invasive transition. Cancer Res, 68, 1378–1387.

86. Lombardi, M.L., Jaalouk, D.E., Shanahan, C.M., Burke, B., Roux, K.J. and Lammerding, J. (2011) The interaction between nesprins and sun proteins at the nuclear envelope is critical for force transmission between the nucleus and cytoskeleton. J Biol Chem, 286, 26743–26753.

87. Swift, J., Ivanovska, I.L., Buxboim, A., Harada, T., Dingal, P.C., Pinter, J., Pajerowski, J.D., Spinler, K.R., Shin, J.W., Tewari, M. et al. (2013) Nuclear lamin-A scales with tissue stiffness and enhances matrix-directed differentiation. Science, 341, 1240104.

88. Kubben, N. and Misteli, T. (2017) Shared molecular and cellular mechanisms of premature ageing and ageing-associated diseases. Nat Rev Mol Cell Biol, 18, 595–609.

89. Gonzalez-Suarez, I., Redwood, A.B., Perkins, S.M., Vermolen, B., Lichtensztejin, D., Grotsky, D.A., Morgado-Palacin, L., Gapud, E.J., Sleckman, B.P., Sullivan, T. et al. (2009) Novel roles for A-type lamins in telomere biology and the DNA damage response pathway. EMBO J, 28, 2414–2427.

90. Gibbs-Seymour, I., Markiewicz, E., Bekker-Jensen, S., Mailand, N. and Hutchison, C.J. (2015) Lamin A/C-dependent interaction with 53BP1 promotes cellular responses to DNA damage. Aging Cell, 14, 162–169.

91. Etourneaud, L., Moussa, A., Rass, E., Genet, D., Willaume, S., Chabance-Okumura, C., Wanschoor, P., Picotto, J., Theze, B., Depagne, J. et al. (2021) Lamin B1 sequesters 53BP1 to control its recruitment to DNA damage. Sci Adv, 7.

92. Todorovski, V., Fox, A.H. and Choi, Y.S. (2020) Matrix stiffness-sensitive long noncoding RNA NEAT1 seeded paraspeckles in cancer cells. Mol Biol Cell, 31, 1654–1662.

93. Dupont, S., Morsut, L., Aragona, M., Enzo, E., Giulitti, S., Cordenonsi, M., Zanconato, F., Le Digabel, J., Forcato, M., Bicciato, S., et al. (2011) Role of YAP/TAZ in mechanotransduction. Nature, 474, 179–183.

94. Elosegui-Artola, A., Andreu, I., Beedle, A.E.M., Lezamiz, A., Uroz, M., Kosmalska, A.J., Oria, R., Kechagia, J.Z., Rico-Lastres, P., Le Roux, A.L., et al. (2017) Force Triggers YAP Nuclear Entry by Regulating Transport across Nuclear Pores. Cell, 171, 1397–1410 e1314.

95. Liu-Chittenden, Y., Huang, B., Shim, J.S., Chen, Q., Lee, S.J., Anders, R.A., Liu, J.O. and Pan, D. (2012) Genetic and pharmacological disruption of the TEAD-YAP complex suppresses the oncogenic activity of YAP. Genes Dev, 26, 1300–1305.

96. Wang, C., Zhu, X., Feng, W., Yu, Y., Jeong, K., Guo, W., Lu, Y. and Mills, G.B. (2016) Verteporfin inhibits YAP function through up-regulating 14-3-3sigma sequestering YAP in the cytoplasm. Am J Cancer Res, 6, 27–37.

97. Monroe, T.O., Hill, M.C., Morikawa, Y., Leach, J.P., Heallen, T., Cao, S., Krijger, P.H.L., de Laat, W., Wehrens, X.H.T., Rodney, G.G., et al. (2019) YAP Partially Reprograms Chromatin Accessibility to Directly Induce Adult Cardiogenesis In Vivo. Dev Cell, 48, 765–779 e767.

98. Fetiva, M.C., Liss, F., Gertzmann, D., Thomas, J., Gantert, B., Vogl, M., Sira, N., Weinstock, G., Kneitz, S., Ade, C.P. et al. (2023) Oncogenic YAP mediates changes in chromatin accessibility and activity that drive cell cycle gene expression and cell migration. Nucleic Acids Res, 51, 4266–4283.

99. Milacic, M., Beavers, D., Conley, P., Gong, C., Gillespie, M., Griss, J., Haw, R., Jassal, B., Matthews, L., May, B. et al. (2024) The Reactome Pathway Knowledgebase 2024. Nucleic Acids Res, 52, D672–D678.

100. Raab, M., Gentili, M., de Belly, H., Thiam, H.R., Vargas, P., Jimenez, A.J., Lautenschlaeger, F., Voituriez, R., Lennon-Dumenil, A.M., Manel, N., et al. (2016) ESCRT III repairs nuclear envelope ruptures during cell migration to limit DNA damage and cell death. Science, 352, 359–362.

101. Denais, C.M., Gilbert, R.M., Isermann, P., McGregor, A.L., te Lindert, M., Weigelin, B., Davidson, P.M., Friedl, P., Wolf, K. and Lammerding, J. (2016) Nuclear envelope rupture and repair during cancer cell migration. Science, 352, 353–358.

102. Xia, Y., Ivanovska, I.L., Zhu, K., Smith, L., Irianto, J., Pfeifer, C.R., Alvey, C.M., Ji, J., Liu, D., Cho, S. et al. (2018) Nuclear rupture at sites of high curvature compromises retention of DNA repair factors. J Cell Biol, 217, 3796–3808.

103. Shah, P., Hobson, C.M., Cheng, S., Colville, M.J., Paszek, M.J., Superfine, R. and Lammerding, J. (2021) Nuclear Deformation Causes DNA Damage by Increasing Replication Stress. Curr Biol, 31, 753–765 e756.

104. Cordes, N., Seidler, J., Durzok, R., Geinitz, H. and Brakebusch, C. (2006) beta1-integrin-mediated signaling essentially contributes to cell survival after radiation-induced genotoxic injury. Oncogene, 25, 1378–1390.

105. Dickreuter, E., Eke, I., Krause, M., Borgmann, K., van Vugt, M.A. and Cordes, N. (2016) Targeting of beta1 integrins impairs DNA repair for radiosensitization of head and neck cancer cells. Oncogene, 35, 1353–1362.

106. Kowalski-Chauvel, A., Modesto, A., Gouaze-Andersson, V., Baricault, L., Gilhodes, J., Delmas, C., Lemarie, A., Toulas, C., Cohen-Jonathan-Moyal, E. and Seva, C. (2018) Alpha-6 integrin promotes radioresistance of glioblastoma by modulating DNA damage response and the transcription factor Zeb1. Cell Death Dis, 9, 872.

107. Fang, H., Ren, W., Cui, Q., Liang, H., Yang, C., Liu, W., Wang, X., Liu, X., Shi, Y., Feng, J. et al. (2023) Integrin beta4 promotes DNA damage-related drug resistance in triple-negative breast cancer via TNFAIP2/IQGAP1/RAC1. Elife, 12.

108. Nair, R.R., Hsu, J., Jacob, J.T., Pineda, C.M., Hobbs, R.P. and Coulombe, P.A. (2021) A role for keratin 17 during DNA damage response and tumor initiation. Proc Natl Acad Sci U S A, 118.

109. Stenvall, C.A., Nystrom, J.H., Butler-Hallissey, C., Jansson, T., Heikkila, T.R.H., Adam, S.A., Foisner, R., Goldman, R.D., Ridge, K.M. and Toivola, D.M. (2022) Cytoplasmic keratins couple with and maintain nuclear envelope integrity in colonic epithelial cells. Mol Biol Cell, 33, ar121.

110. Herrmann, H. and Aebi, U. (2016) Intermediate Filaments: Structure and Assembly. Cold Spring Harb Perspect Biol, 8.

111. Ewald, C.Y. and Nystrom, A. (2023) Mechanotransduction through hemidesmosomes during aging and longevity. J Cell Sci, 136.

112. Deng, M., Lin, J., Nowsheen, S., Liu, T., Zhao, Y., Villalta, P.W., Sicard, D., Tschumperlin, D.J., Lee, S., Kim, J. et al. (2020) Extracellular matrix stiffness determines DNA repair efficiency and cellular sensitivity to genotoxic agents. Sci Adv, 6.

113. Pefani, D.E. and O’Neill, E. (2016) Hippo pathway and protection of genome stability in response to DNA damage. FEBS J, 283, 1392–1403.

114. Chang, H.A., Ou Yang, R.Z., Su, J.M., Nguyen, T.M.H., Sung, J.M., Tang, M.J. and Chiu, W.T. (2023) YAP nuclear translocation induced by HIF-1alpha prevents DNA damage under hypoxic conditions. Cell Death Discov, 9, 385.

115. Shu, Y., Jin, X., Ji, M., Zhang, Z., Wang, X., Liang, H., Lu, S., Dong, S., Lin, Y., Guo, Y. et al. (2024) Ku70 Binding to YAP Alters PARP1 Ubiquitination to Regulate Genome Stability and Tumorigenesis. Cancer Res, 84, 2836–2855.

116. Goodarzi, A.A., Noon, A.T., Deckbar, D., Ziv, Y., Shiloh, Y., Lobrich, M. and Jeggo, P.A. (2008) ATM signaling facilitates repair of DNA double-strand breaks associated with heterochromatin. Mol Cell, 31, 167–177.

117. Smeenk, G. and van Attikum, H. (2013) The chromatin response to DNA breaks: leaving a mark on genome integrity. Annu Rev Biochem, 82, 55–80.

118. Aymard, F., Bugler, B., Schmidt, C.K., Guillou, E., Caron, P., Briois, S., Iacovoni, J.S., Daburon, V., Miller, K.M., Jackson, S.P. et al. (2014) Transcriptionally active chromatin recruits homologous recombination at DNA double-strand breaks. Nat Struct Mol Biol, 21, 366–374.

119. Downs, J.A. and Gasser, S.M. (2024) Chromatin remodeling and spatial concerns in DNA double-strand break repair. Curr Opin Cell Biol, 90, 102405.

120. Killaars, A.R., Grim, J.C., Walker, C.J., Hushka, E.A., Brown, T.E. and Anseth, K.S. (2019) Extended Exposure to Stiff Microenvironments Leads to Persistent Chromatin Remodeling in Human Mesenchymal Stem Cells. Adv Sci (Weinh), 6, 1801483.

121. Alisafaei, F., Jokhun, D.S., Shivashankar, G.V. and Shenoy, V.B. (2019) Regulation of nuclear architecture, mechanics, and nucleocytoplasmic shuttling of epigenetic factors by cell geometric constraints. Proc Natl Acad Sci U S A, 116, 13200–13209.

122. Killaars, A.R., Walker, C.J. and Anseth, K.S. (2020) Nuclear mechanosensing controls MSC osteogenic potential through HDAC epigenetic remodeling. Proc Natl Acad Sci U S A, 117, 21258–21266.

123. Murga, M., Jaco, I., Fan, Y., Soria, R., Martinez-Pastor, B., Cuadrado, M., Yang, S.M., Blasco, M.A., Skoultchi, A.I. and Fernandez-Capetillo, O. (2007) Global chromatin compaction limits the strength of the DNA damage response. J Cell Biol, 178, 1101–1108.

124. Kim, Y.C., Gerlitz, G., Furusawa, T., Catez, F., Nussenzweig, A., Oh, K.S., Kraemer, K.H., Shiloh, Y. and Bustin, M. (2009) Activation of ATM depends on chromatin interactions occurring before induction of DNA damage. Nat Cell Biol, 11, 92–96.

125. Sanz, L.A., Hartono, S.R., Lim, Y.W., Steyaert, S., Rajpurkar, A., Ginno, P.A., Xu, X. and Chedin, F. (2016) Prevalent, Dynamic, and Conserved R-Loop Structures Associate with Specific Epigenomic Signatures in Mammals. Mol Cell, 63, 167–178.

126. Salas-Armenteros, I., Klunder, M., Vermeulen, W. and Tresini, M. (2025) Crosstalk between chromatin state and ATM signalling in DNA damage-induced transcription stress. EMBO J, 44, 5564–5594.

127. Degnim, A.C., Visscher, D.W., Hoskin, T.L., Frost, M.H., Vierkant, R.A., Vachon, C.M., Shane Pankratz, V., Radisky, D.C. and Hartmann, L.C. (2012) Histologic findings in normal breast tissues: comparison to reduction mammaplasty and benign breast disease tissues. Breast Cancer Res Treat, 133, 169–177.

128. Divoux, A., Tordjman, J., Lacasa, D., Veyrie, N., Hugol, D., Aissat, A., Basdevant, A., Guerre-Millo, M., Poitou, C., Zucker, J.D. et al. (2010) Fibrosis in human adipose tissue: composition, distribution, and link with lipid metabolism and fat mass loss. Diabetes, 59, 2817–2825.

129. Gaber, M., Quentel, A., Holmes, J., Lepetit, C., Triki, H., Wilson, A., Payne, V., Tenvooren, I., Dehours, C., Peoples, A. et al. (2025) Obesity increases DNA damage in the breast epithelium. Breast Cancer Res, 27, 11.

130. Chaudhuri, O., Koshy, S.T., Branco da Cunha, C., Shin, J.W., Verbeke, C.S., Allison, K.H. and Mooney, D.J. (2014) Extracellular matrix stiffness and composition jointly regulate the induction of malignant phenotypes in mammary epithelium. Nat Mater, 13, 970–978.

131. Hayward, M.K., Muncie, J.M. and Weaver, V.M. (2021) Tissue mechanics in stem cell fate, development, and cancer. Dev Cell, 56, 1833–1847.

132. Northey, J.J., Hayward, M.K., Yui, Y., Stashko, C., Kai, F., Mouw, J.K., Thakar, D., Lakins, J.N., Ironside, A.J., Samson, S. et al. (2024) Mechanosensitive hormone signaling promotes mammary progenitor expansion and breast cancer risk. Cell Stem Cell, 31, 106–126 e113.

133. Narain, R., Muncie-Vasic, J.M. and Weaver, V.M. (2025) Forcing the code: tension modulates signaling to drive morphogenesis and malignancy. Genes Dev, 39, 163–181.

134. Jones, M., Percival, H., Hales, A., Wood, A., Sun, H., Tennant, F., Broadberry, E., Skaria, E., Barnes, H., Zindy, E. et al. (2025) Matrix stiffness drives alterations in aldehyde metabolism, inducing DNA damage and transformation. Sci Rep, 15, 29127.

135. Tharp, K.M., Higuchi-Sanabria, R., Timblin, G.A., Ford, B., Garzon-Coral, C., Schneider, C., Muncie, J.M., Stashko, C., Daniele, J.R., Moore, A.S. et al. (2021) Adhesion-mediated mechanosignaling forces mitohormesis. Cell Metab, 33, 1322–1341 e1313.

136. DeFilippis, R.A., Chang, H., Dumont, N., Rabban, J.T., Chen, Y.Y., Fontenay, G.V., Berman, H.K., Gauthier, M.L., Zhao, J., Hu, D. et al. (2012) CD36 repression activates a multicellular stromal program shared by high mammographic density and tumor tissues. Cancer Discov, 2, 826–839.

137. Fan, Y., An, C., Wang, Z., Luo, J., Wang, W., Luo, Q. and Song, G. (2025) Matrix stiffening induces hepatocyte functional impairment and DNA damage via the Piezo1‒ERK1/2 signaling pathway. J Physiol Biochem.

138. Wu, H.C., Kehm, R., Santella, R.M., Brenner, D.J. and Terry, M.B. (2022) DNA repair phenotype and cancer risk: a systematic review and meta-analysis of 55 case-control studies. Sci Rep, 12, 3405.

