## Supplementary material for "Mechanotransduction tuned by intermediate filaments promotes DNA repair in stiff environments": SI

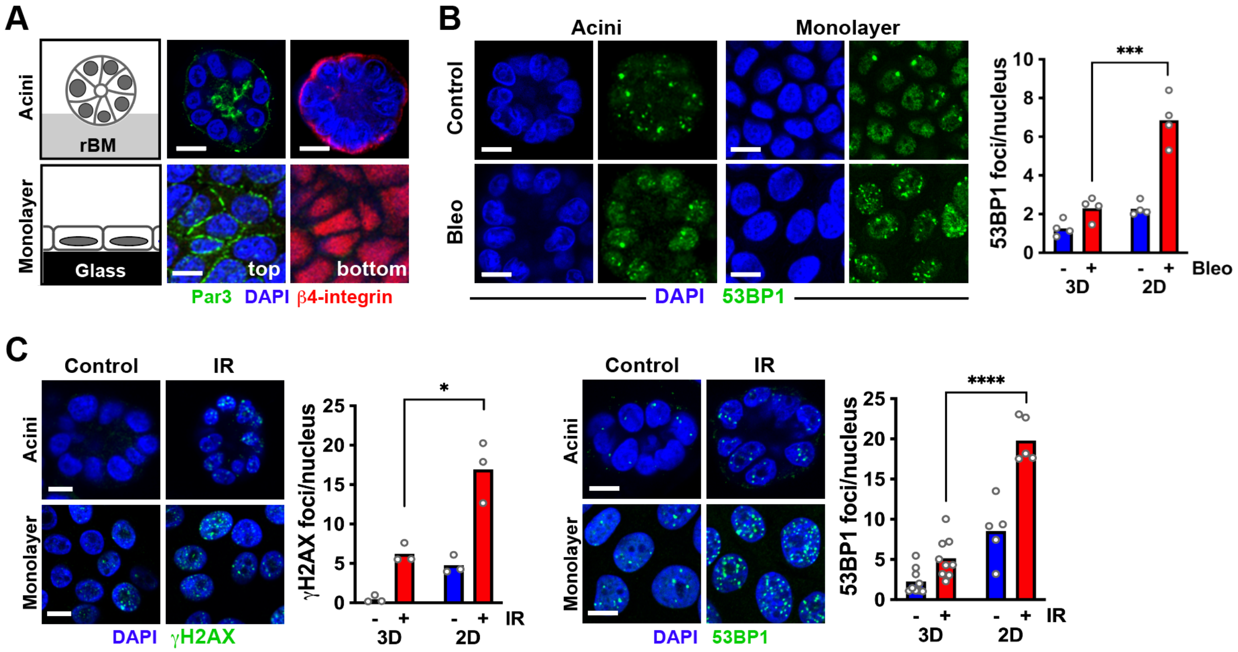


**Supplementary Figure S1.** Breast acini and cell monolayers respond differently to DNA damage. (**A**) Differentiation of S1 cells on soft reconstituted basement membrane (rBM) hydrogel or on glass, resulting in acini and monolayers, respectively. Cell differentiation is illustrated by the proper localization of Par3 and β4-integrin apical-basal polarity markers. (**B**) 53BP1 foci in S1 acini and monolayers after bleomycin treatment (Bleo; 20 mU/ml; 2h). ***, $P < 0.001$ (unpaired t-test). (**C**) Formation of γH2AX and 53BP1 repair foci in S1 acini and monolayers in response to ionizing radiation (IR; 3 Gy). Controls were untreated. *, $P < 0.05$ and ****, $P < 0.0001$ (unpaired t-test). Scale bars, 10 μm.

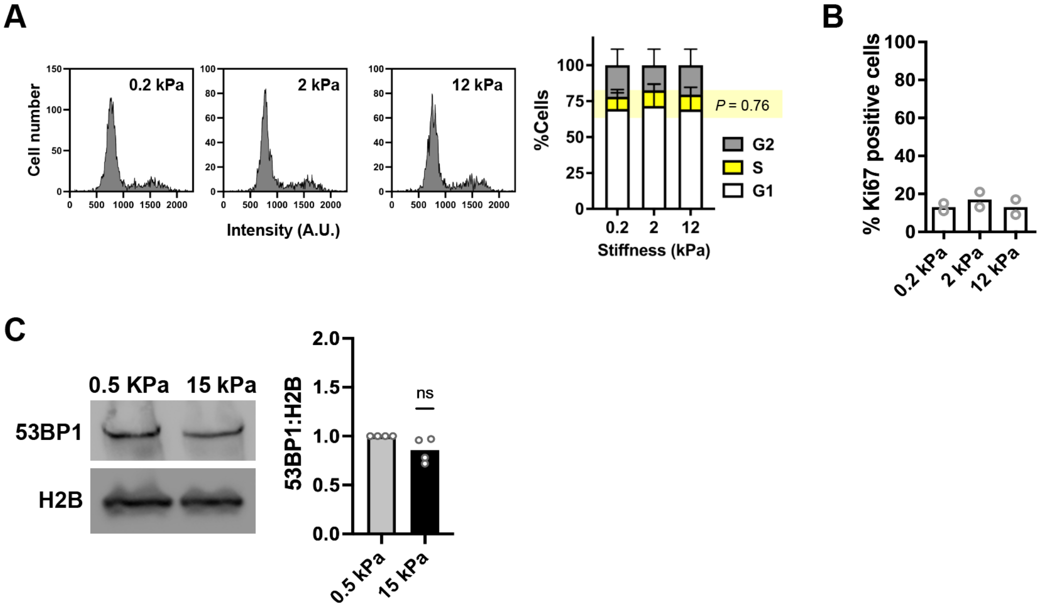


**Supplementary Figure S2.** Phenotype of S1 cells in different stiffness conditions. (**A**) Cell cycle distribution on PA gels with different stiffnesses (N = 3; mean ± SD). Frequency distribution plots of DAPI intensities determined by imaging cytometry are shown for illustration. Statistical analysis of S-phase content by ANOVA. (**B**) Proportion of Ki67-positive S1 cells on the different stiffnesses. (**C**) Western blot analysis of 53BP1 expression in cells cultured on soft and stiff silicone substrates. Ns, not significant (one-sample t-test).

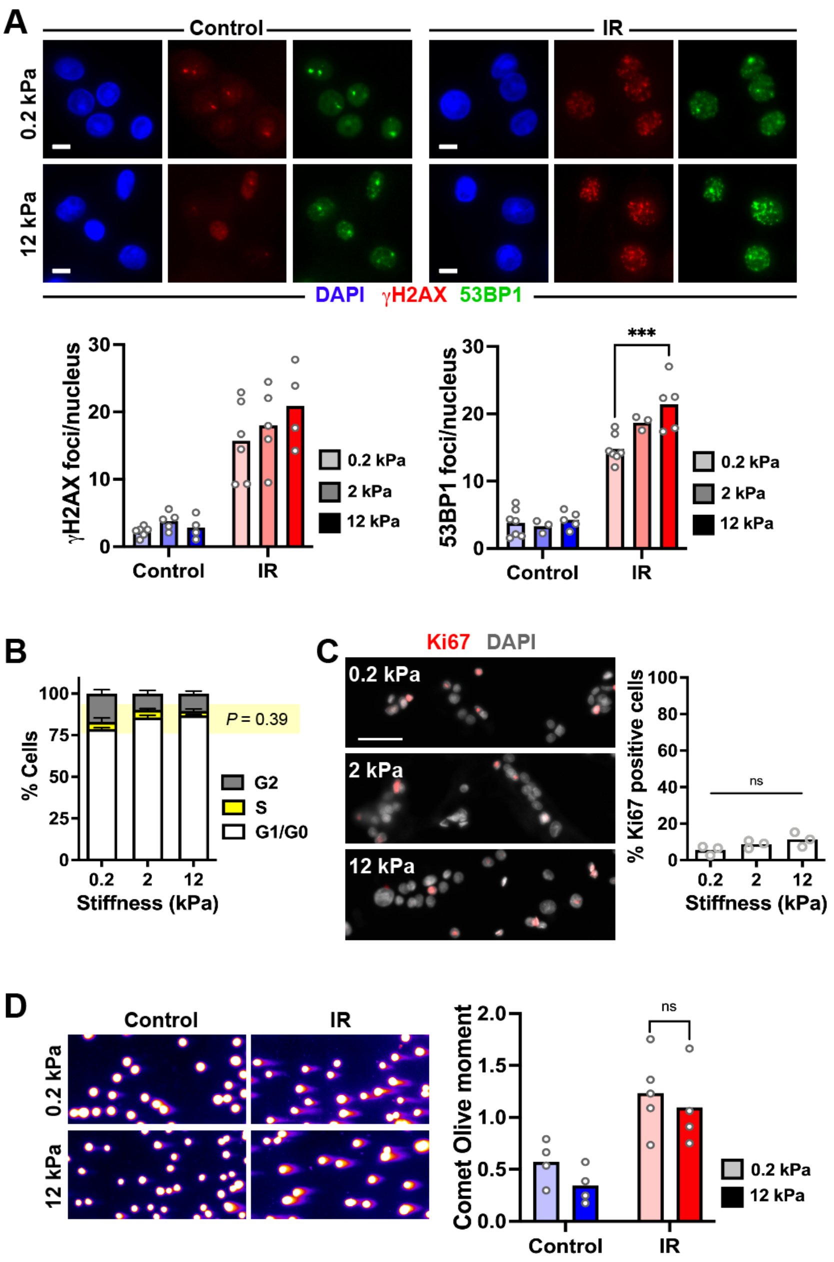


**Supplementary Figure S3.** Substrate stiffness exacerbates DSB repair foci accumulation in non-immortalized breast epithelial cells. (**A**) γH2AX and 53BP1 repair foci in human mammary epithelial cells (HMEC) cultured on polyacrylamide gels with different stiffnesses. Cells were either untreated (control) or irradiated (IR, 3 Gy). ***, $P$ = 0.0001 (ANOVA and Bonferroni's multiple comparison test). (**B**) Cell cycle distribution of HMEC cells cultured on different stiffnesses. Data represent mean ± SD from 3 experiments. (**C**) Proportion of Ki67-positive HMEC cells in the different stiffness conditions. Ns, not significant (unpaired t-test in B; ANOVA in C and D). Scale bar, 50 µm. (**D**) Quantification of DSBs by neutral comet assay in irradiated HMEC cells (directly after IR) and in nonirradiated controls. Ns, not significant (unpaired t-test).

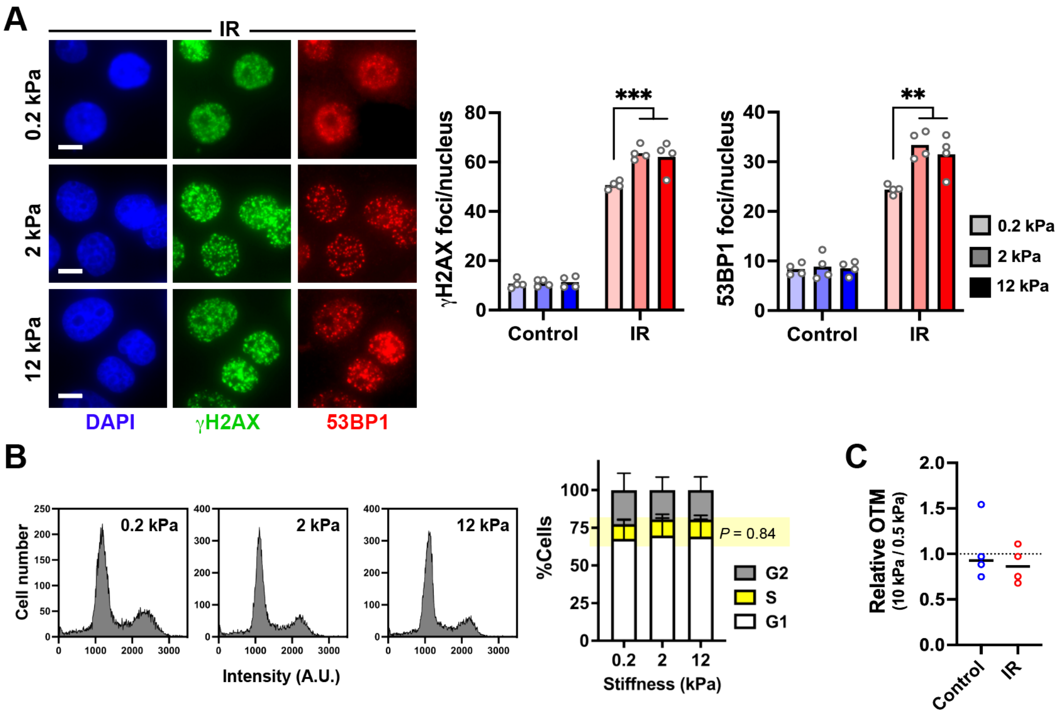


**Supplementary Figure S4.** Radiation response of breast cancer cells depends on the stiffness of the substrate. (**A**) Immunostaining for γH2AX and 53BP1 of HMT-3522 T4 cells on polyacrylamide gels with different elastic moduli. Cells were either untreated (control) or irradiated (IR) and left to recover for 1h before immunostaining. Scale bars, 10 µm. **, $P < 0.01$ and ***, $P < 0.001$ (ANOVA and Bonferroni's multiple comparison test). (**B**) Cell cycle distribution by imaging cytometry (N = 3; mean ± SD). Statistical analysis of S-phase content by ANOVA. (**C**) Quantification of DSBs by neutral comet assay. Relative Olive tail moment (OTM) values are shown.

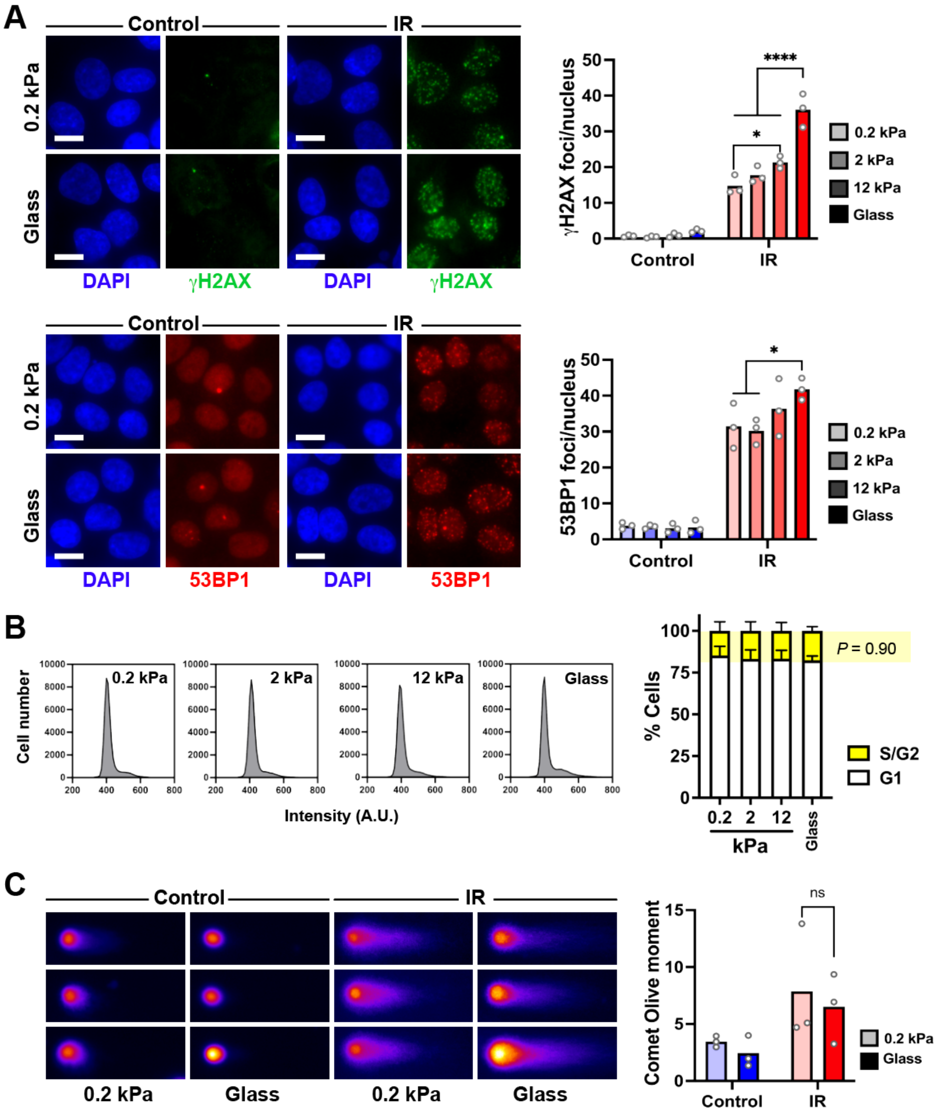


**Supplementary Figure S5.** Differential radiation responses of liver epithelial cells on soft and stiff substrates. (**A**) Immunostaining for γH2AX and 53BP1 of HepaRG cells cultured on polyacrylamide gels with different elastic moduli, as well as on glass. Cells were either untreated (control) or irradiated (IR) and left to recover for 1h. Scale bars, 10 µm. *, $P < 0.05$ and ****, $P < 0.0001$ (ANOVA and Bonferroni's multiple comparison test). (**B**) Cell cycle distribution by imaging cytometry (N = 3; mean ± SD). Statistical analysis of S-phase content by ANOVA. (**C**) Quantification of DSBs by neutral comet assay. Ns, not significant (unpaired t-test).

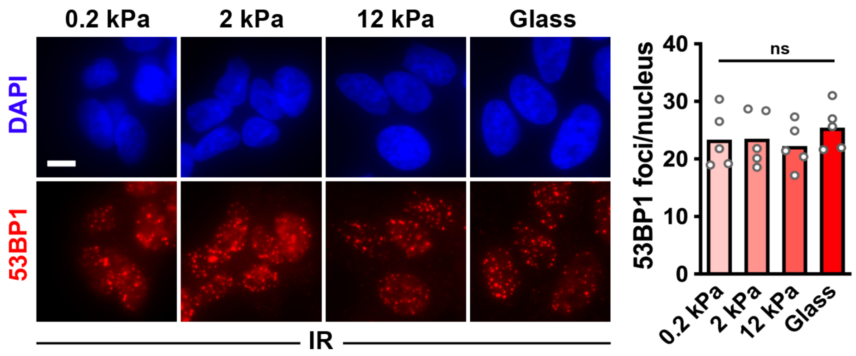


**Supplementary Figure S6.** Substrate stiffness does not influence the DNA damage response of osteosarcoma cells. 53BP1 repair foci in U2OS cells cultured on glass and on polyacrylamide gels with different elastic moduli, 1 h after irradiation (IR). Scale bar, 10 µm. Ns, not significant (ANOVA and Bonferroni's multiple comparison test).

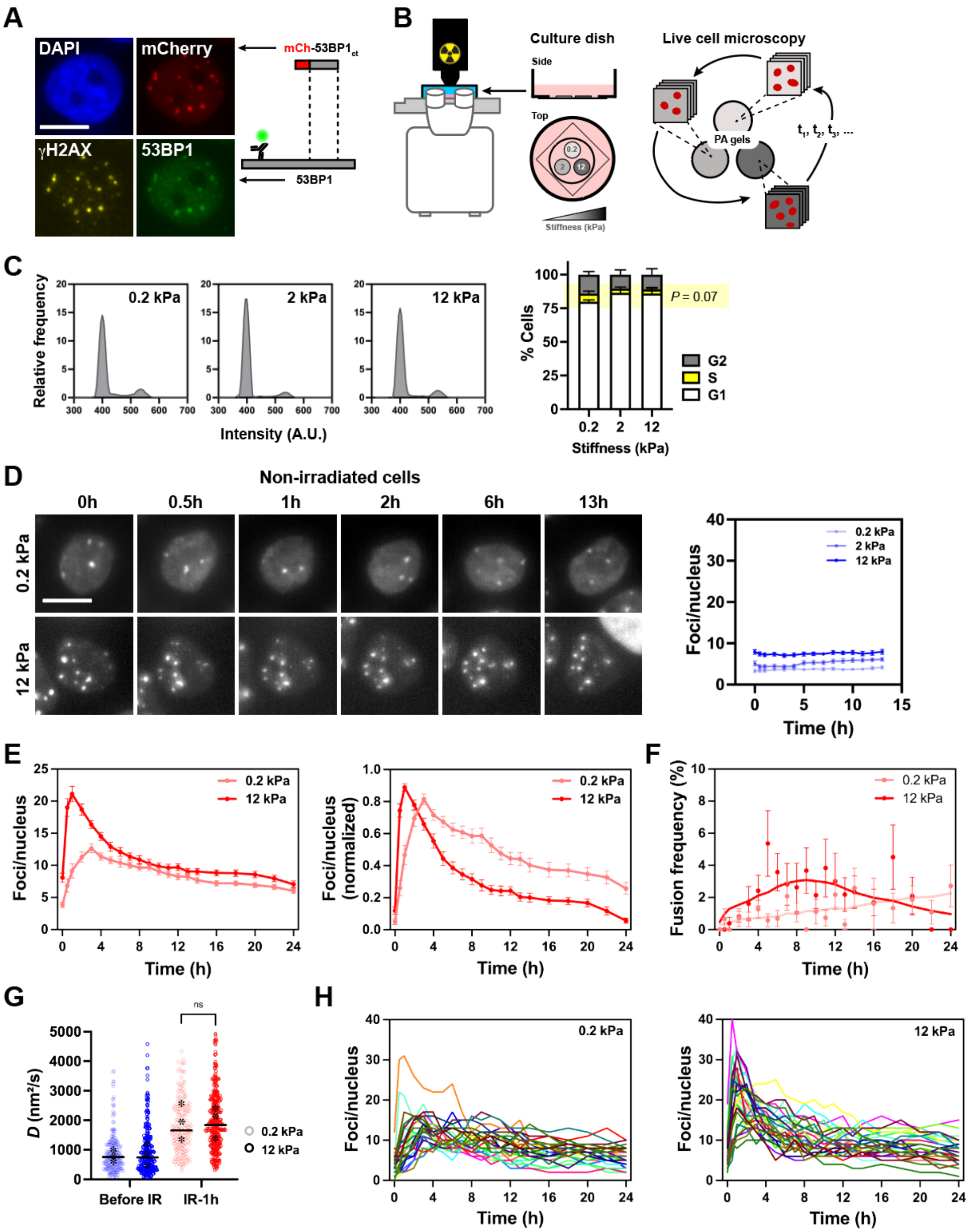


**Supplementary Figure S7.** Analysis of the DNA damage response in live cells. (**A**) MCF10A cells expressing mCherry-53BP1ct, immunostained for detection of γH2AX and endogenous 53BP1. Bleomycin (20 mU/ml; 1h) was used to induce DSBs. Scale bar, 10 µm. (**B**) Cell culture dish with thin polyacrylamide gel disks for imaging in rapid succession cells on different stiffnesses. (**C**) Cell cycle distribution of MCF10A-mCh-53BP1ct cells (N = 3; mean ± SD). Statistical analysis of S-phase content by ANOVA. (**D**) Time-lapse imaging of non-irradiated MCF10A cells expressing mCherry-53BP1ct, on soft or stiff substrates. DSB repair foci are quantified in the graph (N = 25-30 cells per condition). (**E**) 24h time-course of mCherry-53BP1ct foci after irradiation (N = 30 cells per condition). For normalized IR responses (right), minimal and maximal foci numbers were set to 0 and 1, respectively, for each cell. (**F**) Frequency of mCh-53BP1$_{ct}$ foci fusions averaged per cell at each time point. (**G**) Diffusion coefficient of mCh-53BP1$_{ct}$ foci before and after irradiation on soft or stiff substrates. Ns, not significant (Mann-Whithney; N > 180 cells from 3 replicates). Average values from each replicate are show with *. (**H**) Foci count for individual cells on soft and stiff substrates.

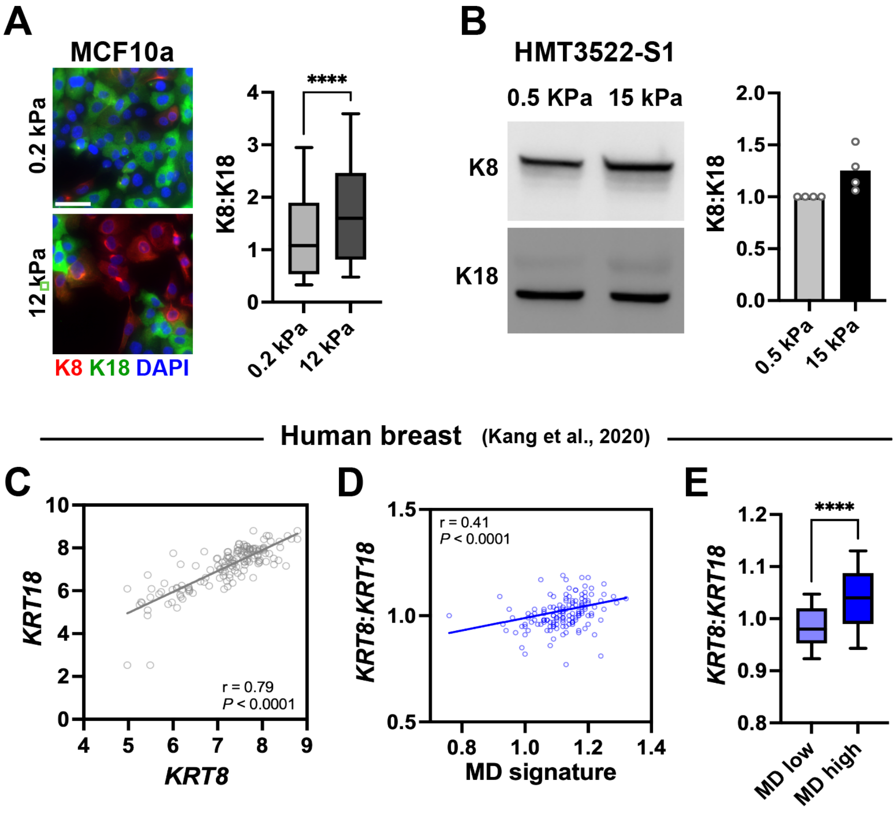


**Supplementary Figure S8.** Cytokeratin expression in soft and stiff contexts. (**A**) Relative levels of K8 and K18 in MCF10A cells cultured on soft and stiff polyacrylamide gels, detected by immunostaining. Scale bar, 50 µm. ****, $P < 0.0001$ (Mann-Whitney test; N = 500-1000 cells from 2 biological replicates). (**B**) Western blot analysis of relative K8 and K18 levels in HMT-3522 S1 cells grown on soft and stiff silicone substrates. (**C-E**) Analysis of normal breast tissue transcriptomic data. Batch-corrected RNAseq values were retrieved using UCSC Xena (https://xenabrowser.net/). The correlation between *KRT8* and *KRT18* gene expression (**C**), as well as the correlation between the *KRT8*:*KRT18* expression ratio and a MD signature (**D**), was evaluated using Spearman's correlation coefficient (r). (**E**) *KRT8:KRT18* gene expression ratio in 'high-' and 'low-MD' tissues (based on the median of MD signature scores). ****, $P < 0.0001$ (Mann-Whitney test; N=72/72). Genes used for the MD signature include *COL1A1*, *COL1A2, COL5A1*, *COL12A1*, *COL16A1*, *LUM*, *BGN*, *ECM1*, *MGP*, *DPT*, *POSTN*, and *DCN*, with normalized expression values averaged.

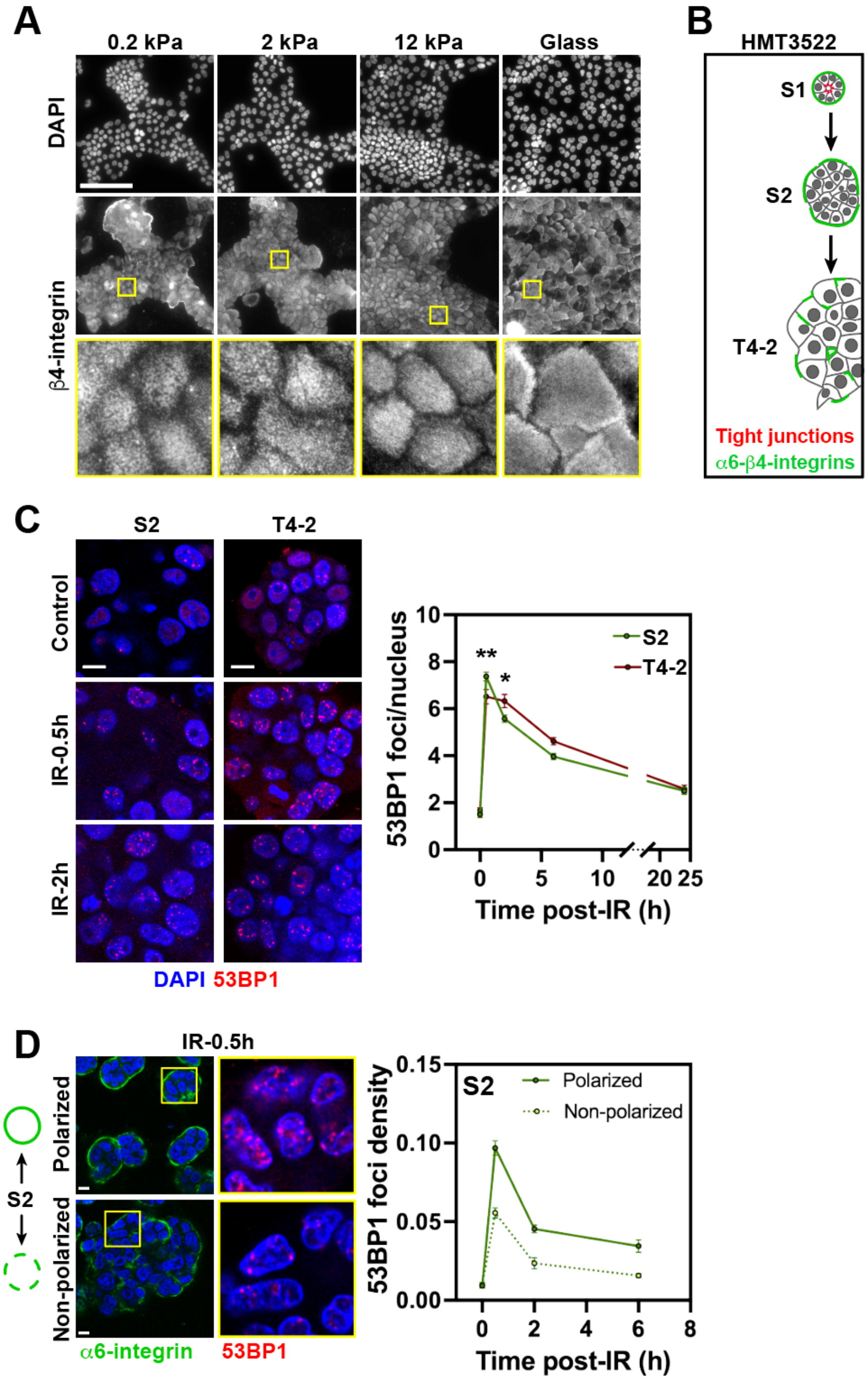


**Supplementary Figure S9.** Integrin mechanotransduction in the DDR. (**A**) Immunostaining for β4-integrin in HMT-3522 S1 cells on substrates with different stiffnesses. (**B**) Illustration of apicobasal polarity loss in the HMT-3522 breast cancer progression series. (**C**) 53BP1 repair foci in nodules from HMT-3522 S2 and HMT-3522 T4-2 cells in response to irradiation (IR; 3 Gy). **, $P = 0.006$ and *, $P = 0.049$ (ANOVA and Šidák's multiple comparison test; N = 10-75 nodules from 4 biological replicates). (**D**) Accumulation of 53BP1 foci in S2 nodules according to the polarization status of their hemidesmosomes (N > 70 nuclei). Scale bars, 100 µm (A) and 10 µm (B-D).

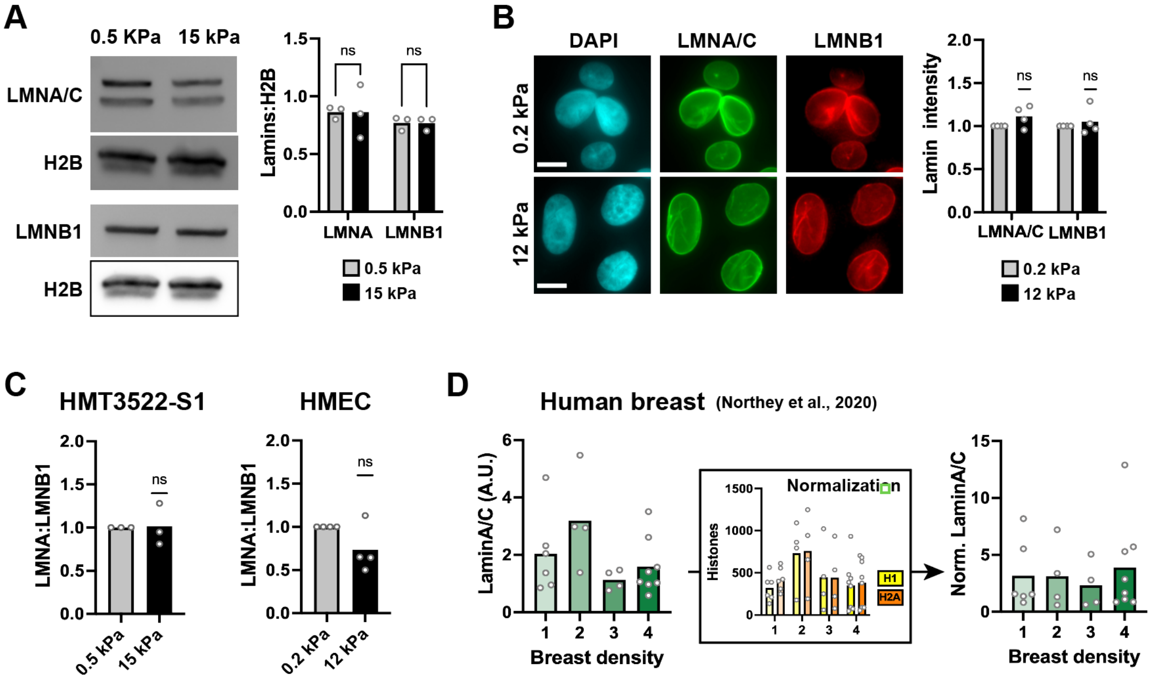


**Supplementary Figure S10.** Lamins are not differentially expressed in breast epithelial cells on soft vs. stiff substrates. (**A**) Western blot analysis of lamins levels in HMT-3522 S1 cells grown on soft and stiff silicone substrates. Ns, not significant (t-test). (**B**) Quantitative immunostaining analysis (integrated signal intensities) of lamin expression in S1 cells cultured on soft and stiff PA gels. Scale bar, 10 µm. Ns, not significant (one-sample t-test). (**C**) Relative levels of laminA/C (LMNA) and lamin B1 in S1 and HMEC cells, measured by western blot. Ns, not significant (one-sample t-test). (**D**) Lamin A/C protein levels in normal breast tissue with different BIRADS density categories determined by quantitative proteomics by Northey et al. (2020). Data are normalized according to histone levels to account for differences in tissue cellularity.

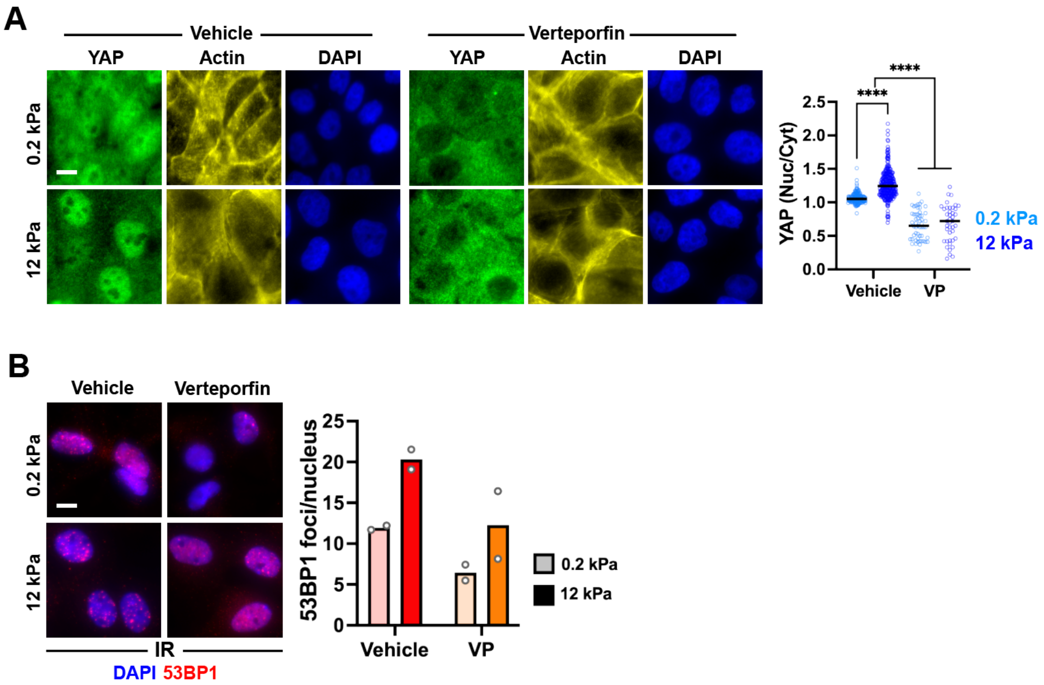


**Supplementary Figure S11.** YAP mediates a mechano-dependent DNA damage response. (**A**) Localization of YAP in MCF10A cells on soft (0.2 kPa) and stiff (12 kPa) PA gels following treatment with verteporfin (VP; 1µM, 6h) or vehicle as control. ****, $P < 0.0001$ (Kruskal-Wallis and Dunn's multiple comparison test N = 40-200 cells). (**B**) IR-induced 53BP1 foci in cells treated with vehicle or VP. Scale bars, 10 µm.

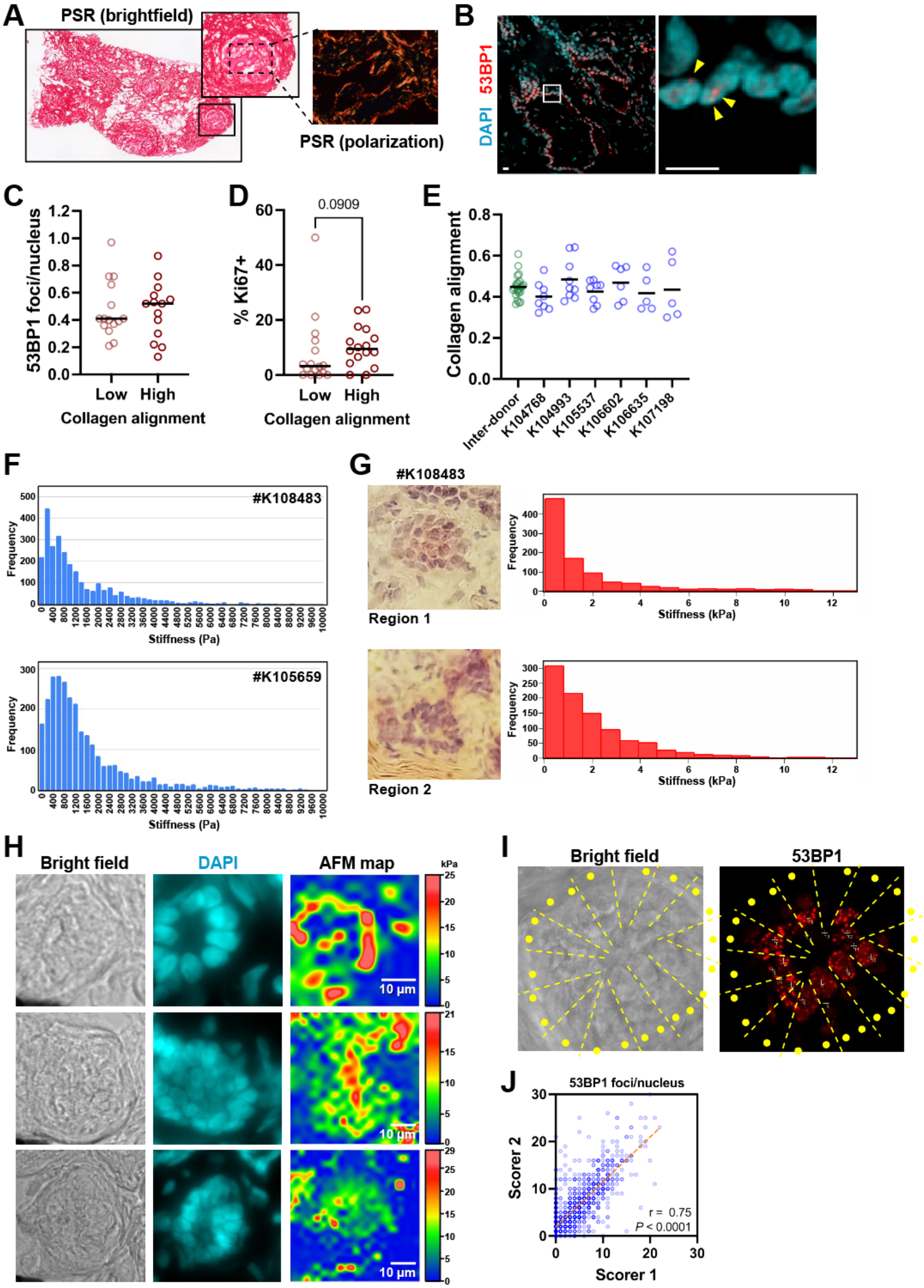

A
PSR (brightfield)
PSR (polarization)
B
DAPI 53BP1
C
53BP1 foci/nucleus
Low
High
Collagen alignment
D
0.0909
% Ki67+
Low
High
Collagen alignment
E
Collagen alignment
Inter-donor
K104768
K104993
K105537
K106602
K106635
K107198
F
#K108483
Frequency
Stiffness (Pa)
#K105659
Frequency
Stiffness (Pa)
G
#K108483
Region 1
Frequency
Stiffness (kPa)
Region 2
Frequency
Stiffness (kPa)
H
Bright field
DAPI
AFM map
kPa
10 µm
I
Bright field
53BP1
J
53BP1 foci/nucleus
Scorer 2
Scorer 1
r = 0.75
P < 0.0001

**Supplementary Figure S12.** Impact of collagen alignment and rheological characteristics of human breast tissue on the DDR. (**A**) Illustration of picrosirius red (PSR) staining of normal breast tissue sections. The inset shows birefringence polarization imaging of an epithelial region to determine collagen alignment. (**B**) Confocal images of 53BP1 immunostaining of a breast tissue section. Scale bars, 10 μm. (**C**) 53BP1 foci and (**D**) the proportion of Ki67-positive cells in breast samples with low vs. high collagen alignment (based on median). Each dot represents a different donor. (**E**) Inter- and intra-individual heterogeneity in collagen alignment. Green dots represent donor averages. Blue dots represent distinct epithelial regions within a tissue, with each value being the average of $N \geq 5$ measurements. (**F**) Distribution of stiffness values from AFM measurements of breast tissue from two donors. (**G**) Stiffness value distribution at different locations within one breast tissue section. (**H**) Local mapping of breast tissue stiffness by AFM. Three regions from the same tissue section are shown. (**I**) Illustration of 53BP1 immunostaining matched with AFM point measurements (yellow dots) in the ECM flanking the epithelial structure. (**J**) Concordance of visual scoring of 53BP1 damage foci between two investigators. r, Spearman correlation.

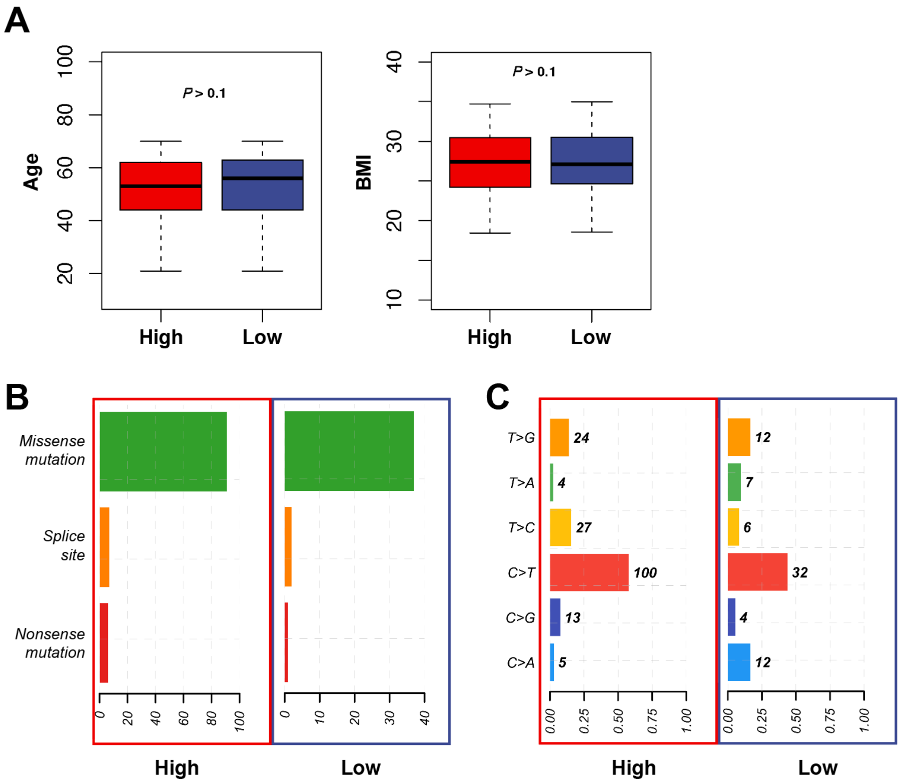


**Supplementary Figure S13.** Mutational analyses of GTEx RNA-Seq data. (**A**) Age and BMI from breast tissue donors, according to the breast density category ('high' and 'low') assigned with a gene expression signature. (**B**) Classification of variants and (**C**) SNV class in the 'high MD' and 'low MD' groups.

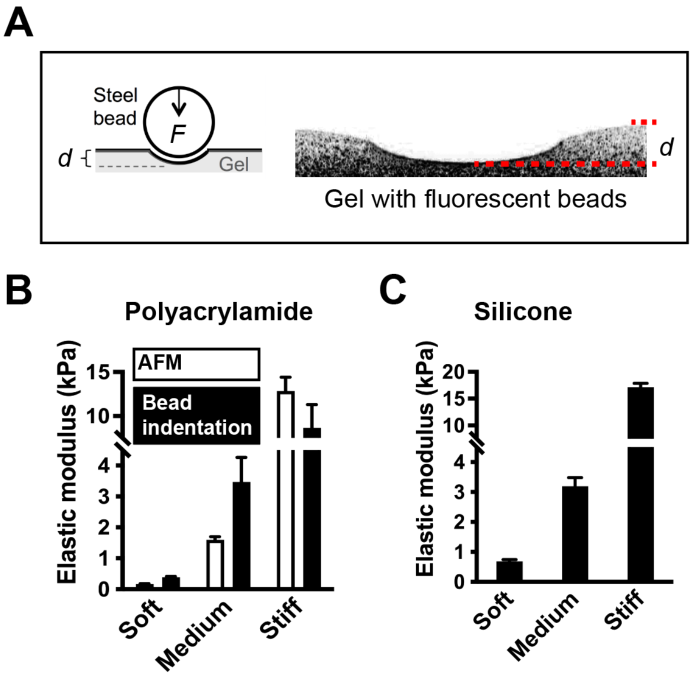


**Supplementary Figure S14.** Rheological characterization of growth substrates. (**A**) Illustration of bead indentation measurements. The indentation distance (*d*) is measured in orthogonal (*x*/*z*) projections of confocal *z*-stacks or by determining the focal *z* position of fluorescent beads at the surface of the substrate with wide field microscopy. (**B**) Microbead indentation and atomic force microscopy (AFM) measurements of polyacrylamide gel substrates (N = 3). (**C**) Bead rheology to verify the elastic modulus of silicone-based cell culture substrates.

**Table S1.** Reactome pathways enriched in low-expressed genes from the high MD group

| ID | Description | Enrichment score | *P* value |
|---|---|---|---|
| R-HSA-381340 | Transcriptional regulation of white adipocyte differentiation | -0.78 | 9.0E-07 |
| R-HSA-194315 | Signaling by Rho GTPases | 0.40 | 3.0E-03 |
| R-HSA-556833 | Metabolism of lipids | -0.32 | 4.8E-03 |
| R-HSA-8979227 | Triglyceride metabolism | -0.56 | 4.9E-03 |
| R-HSA-202733 | Cell surface interactions at the vascular wall | 0.33 | 6.4E-03 |
| R-HSA-1266738 | Developmental Biology | -0.34 | 8.2E-03 |

**Table S2.** Reactome pathways enriched in over-expressed genes from the high MD group

| ID | Description | Enrichment score | *P* value |
|---|---|---|---|
| R-HSA-6809371 | Formation of the cornified envelope | 0.62 | 2.21E-06 |
| R-HSA-6805567 | Keratinization | 0.60 | 5.77E-06 |
| R-HSA-1500931 | Cell-Cell communication | 0.57 | 1.24E-04 |
| R-HSA-446728 | Cell junction organization | 0.59 | 1.29E-04 |
| R-HSA-421270 | Cell-cell junction organization | 0.57 | 3.07E-03 |

**Supplemental movies**

**Movie S1:** Time course of mCh-53BP1$_{ct}$ foci in non-irradiated MCF10A cells

**Movie S2:** Time course of mCh-53BP1$_{ct}$ foci in irradiated MCF10A cells (0.2 kPa)

**Movie S3:** Time course of mCh-53BP1$_{ct}$ foci in irradiated MCF10A cells (12 kPa)

**Movies S4:** MCF10A cell with spinning nucleus

The movies represent 13h of imaging. For irradiation time courses, cells were irradiated after 5 min (second frame of the movies).
